# Crosstalk between RNA m^6^A and DNA methylation regulates transposable element chromatin activation and cell fate in human pluripotent stem cells

**DOI:** 10.1101/2022.09.08.507172

**Authors:** Tongyu Sun, Yueyuan Xu, Yu Xiang, Erik J Soderblom, Yarui Diao

## Abstract

Transposable elements (TEs) are parasitic DNA sequences accounting for over half of the human genome. Tight control of the repression and activation states of TEs is critical for genome integrity, development, immunity, and diseases, including cancer. However, precisely how this regulation is achieved remains unclear. To address this question, we develop a targeted proteomic proximity labeling approach to capture TE-associated proteins in human embryonic stem cells (hESCs). We find that the RNA N6-methyladenosine(m^6^A)-reader, YTHDC2, occupies genomic loci of the primate-specific TE, LTR7/HERV-H, specifically through its interaction with m^6^A-modified HERV-H RNAs. Unexpectedly, YTHDC2 recruits the DNA 5-methylcytosine(5mC)-demethylase, TET1, to remove 5mC from LTR7/HERV-H and prevent epigenetic silencing. Functionally, the YTHDC2/LTR7-axis inhibits neural differentiation of hESCs. Our results reveal both an underappreciated crosstalk between RNA m^6^A and DNA 5mC, the most abundant regulatory modifications of RNA and DNA in eukaryotes, and the fact that in hESCs this interplay controls TE activity and cell fate.

## INTRODUCTION

Transposable elements (TEs) are mobile DNA elements derived from ancient viral infections ^1–4^. Remarkably, in humans, TEs and their remnants account for more than half of our genomic sequences. The activation of TEs often threatens host genome integrity and results in deleterious consequences such as mutations and chromosome rearrangements ^1–4^. Therefore, numerous host defense mechanisms have evolved to suppress TE activation and transcription. In eukaryotes, this silencing is mainly achieved by repressive heterochromatic marks including hypermethylation of DNA 5-methylcytosine (5mC) ^5, 6^ and histone H3K9me3 and H4K20me3 modifications ^7–9^. Most TEs are silenced in the mammalian genome by the host cells. On the other hand, through evolution, some TE sequences have been co-opted as *cis*-regulatory elements (cREs) of the host genome^10^ and control spatial-temporal gene in numerous biological processes ^11^, including embryogenesis, brain development, immune response, and genome organization ^1–4^. In the case of these TEs, a tight balance between the transcriptionally silenced and activated chromatin states is critical for genome integrity and spatial-temporal gene regulation. However, how this tight regulation is achieved is still not fully elucidated. This critical knowledge gap essentially obscures our understanding of the function and regulation of a large proportion of the human genome.

The hypermethylation of TE DNA sequences is a common strategy utilized by host cells to silence TEs among eukaryotes^5, 6^. In mammals, the most common DNA methylation, 5-methylcytosine (5mC) of CpG dinucleotides, is deposited by the *de novo* DNA methyltransferases, DNMT3A and DNMT3B, and maintained by DNMT1^12, 13^. The family of Ten-eleven translocation (TET) methylcytosine dioxygenase, including TET1-3, can remove DNA methylation through the stepwise oxidation of 5mC to 5-hydroxymethycytosine (5hmC), 5-formylcytosine (5fC) and 5-carboxylcytosine (5caC)^14, 15^. In addition to DNA methylation, recently, several studies have revealed a critical and previously unappreciated role for the important process of N6-methyladenosine (m^6^A) of RNA transcripts in regulating TE silencing through histone modifications^16–22^. In eukaryotes, m^6^A is the most abundant form of posttranscriptional RNA modification, regulating numerous biological processes and disease progression^22^. RNA m^6^A is installed by the methyltransferase complex, can be removed by “erasers”, and is recognized by the “reader” proteins, including YTHDC1-2 and YTHDF1-3 that contain a highly conserved m^6^A-binding, YTH-domain^22, 23^. Recent studies show that m^6^A-modified “chromatin-associated regulatory RNAs” (carRNAs), including RNAs transcribed from promoters, enhancers, and TE loci, are bound by the m^6^A reader, YTHDC1, which either induces degradation of TE transcripts or recruits a variety of histone modifiers to induce epigenetic activation or silencing TE genomic loci by modulating histone modifications^24–26^.

While these findings on the cross-talk between RNA m^6^A and histone codes are exciting, the question remains open as to whether the regulatory RNAs can also interplay with the DNA methylation machinery in mammals, another fundamental epigenetic mechanism regulating transcription and chromatin state of regulatory genomic sequences, including TE^27, 28^. Interestingly, in plants, the RNA-directed DNA methylation (RdDM) pathway has been identified for decades, in which non-coding RNAs can direct the addition of DNA methylation to specific DNA sequences, including TE loci^29, 30^. To date, however, the RdDM pathway appears unique to plants. Whether and how regulatory RNA molecules, and/or their epitranscriptome modifications interplay with DNA methylation machinery in other kingdoms remains an intriguing and unanswered question. This critical knowledge gap prevents our comprehensive understanding of the crosstalk between epitranscriptome and epigenome in gene regulation.

Here, we unveil an underappreciated mechanism based on the interplay between RNA m^6^A and DNA methylation in controlling chromatin states and transcription of TE sequences. We develop CARGO-BioID, a CRISPR-based TE-centric proteomics approach, to identify the proteins associated with the genomic loci of the primate-specific TE LTR7/HERV-H (**Fig. 1a**) in human embryonic stem cells (hESCs). We find that the RNA m^6^A-reader, YTHDC2^31^, is recruited to the LTR7 DNA regions through its interaction with m^6^A-modified HERV-H transcripts. Surprisingly, YTHDC2 further recruits the DNA 5mC demethylase TET1^15^ to prevent the silencing of LTR7/HERV-H chromatin via DNA demethylation. Functionally, we show that the YTHDC2/LTR7-axis plays a causal role in preventing the neuronal fate of hESCs. Together, our results establish both new mechanisms controlling TE activity and hESC fate, and crosstalk between RNA m^6^A and regulation of DNA methylation through physical and functional interaction between YTHDC2 and TET1. These findings have the potential to advance our understanding of gene regulation in developmental disorders and many human diseases that involve the regulation of RNA m^6^A and DNA methylation.

**Figure 1.**
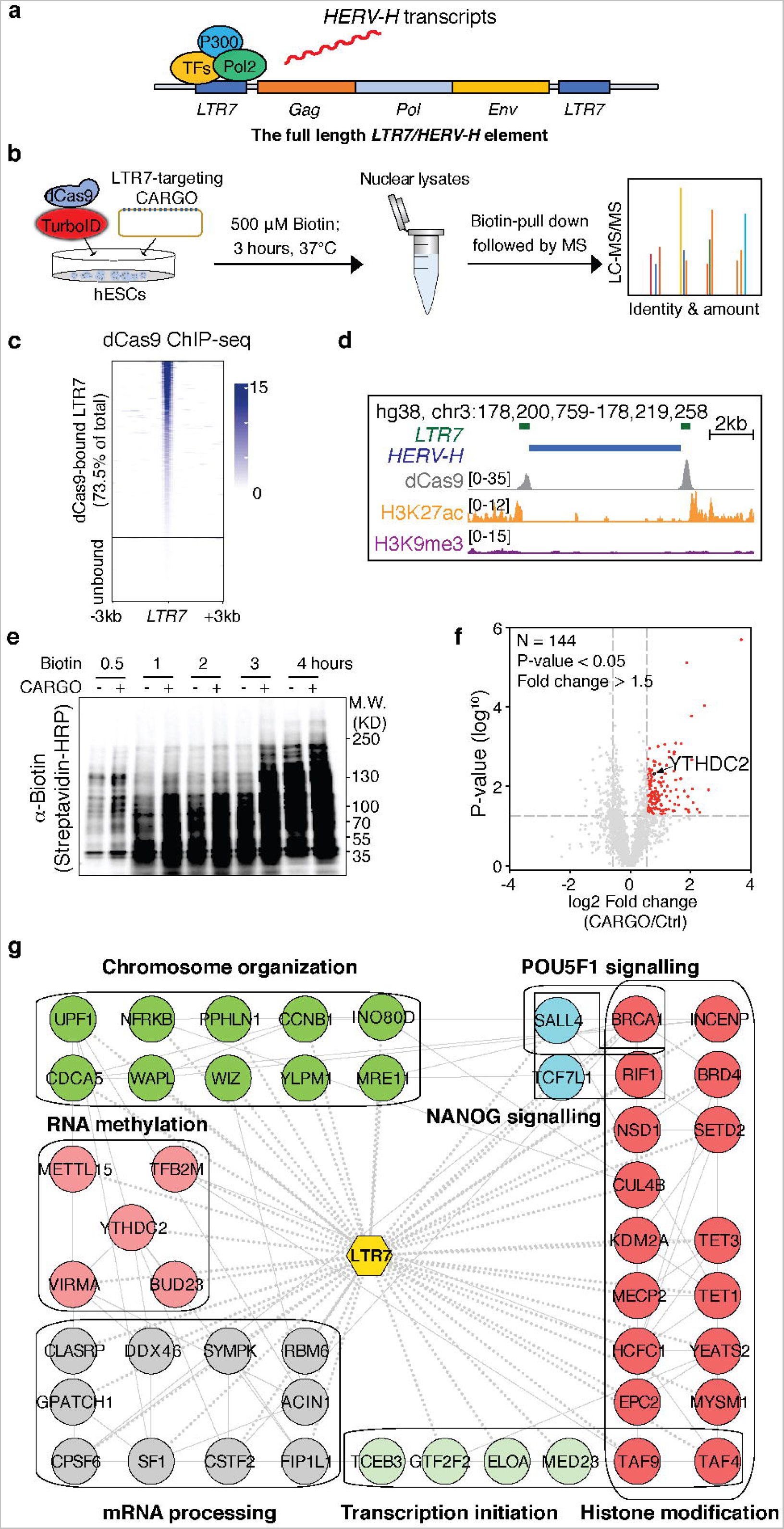
Development of CARGO-BioID to identify LTR7 associated proteome in hESCs. **a,** Schematic of a typical LTR7/HERV-H element. **b,** Illustration of the CARGO-BioID experimental design. **c, d,** Heatmap (**c**) and UCSC genome browser snapshot (**d**) illustrating dCas9 binding, histone H3K27ac and H3K9me3 modifications on a typical LTR7/HERV-H element. **e,** Patterns of nuclear protein biotinylation at indicated experimental conditions. Three hours of biotin labeling is selected for proteomic analysis. **f,** Volcano plot showing 144 significantly enriched (fold change > 1.5, P value < 0.05) proteins that are spatially proximal to the LTR7 loci. YTHDC2 is highlighted in red. **g,** Interaction network and enriched GO and pathway terms of selected LTR7 bound proteins identified in H1 hESC cells. Known protein-protein interaction information is retrieved from the STRING protein interaction database with default confidence cutoff using Cytoscape. Proteins are grouped by their known molecular function in different colors. Node titles show the corresponding gene symbols. Dashed edges show the interactions with LTR7s. Solid edges show the known interactions.

## RESULTS

### CARGO-BioID identifies the transposable elements (TEs)-associated proteome

TE loci have large copy numbers and highly repetitive sequences, as well as locus-specific mutations or indels that accumulated through evolution. Therefore, identification of the effectors controlling the activity of endogenous TE sequences remains technically challenging. To explore potential binding complexes at TEs, we leveraged the multiplexed CRISPR targeting approach “Chimeric Array of gRNA Oligos” (CARGO)^32, 33^ with biotin proximity labeling and developed a CRISPR-based TE-centric proteomics approach, which we termed CARGO-BioID (**Fig. 1b**). In CARGO-BioID, the CARGO construct^32, 33^ enables the simultaneous expression of 15 sgRNAs (**Supplementary Table 1,** Oligo sequences used in this study) to target the conserved sequences of TE sequences in the native chromatin environment^32, 33^ (**Extended Data Fig. 1a**). We fused dCas9 (the catalytic inactive “dead” Cas9 that can bind to but not cleave DNA) with an engineered biotin ligase TurboID^34, 35^, that is predicated to biotinylated proteins proximate to dCas9-TurboID and the DNA sequences to which it is directed^36–39^. The proteins are then purified by streptavidin selection for liquid chromatography-tandem mass spectrometry (LC-MS/MS) analysis (**Fig. 1b** and **Extended Data Fig. 1b, 1c**). By comparing the biotinylated proteins purified from the dCas9-TurboID-expressing cells with CARGO versus those expressing the control non-targeting sgRNAs, we expect to identify proteins that are enriched at TE loci.

We applied CARGO-BioID to the primate-specific TE LTR7/HERV-H (**Fig. 1a**) to study the regulation of TE transcription and chromatin state in the human genome. Epigenome analysis revealed that LTR7/HERV-H were enriched with active histone marks such as histone H3K4me1, H3K4me3, and H3K27ac in pluripotent hESCs (**Extended Data Fig. 1d**), with less representation of these marks in more differentiated lineages like mesendoderm, neural progenitors, and trophoblast cells^40, 41^ (**Extended Data Fig. 1e**). LTR7 sequences have been found to be bound by transcription factors and co-factors in hESC and iPSC lines, which drive the expression of HERV-H transcripts and the retroviral genes *Gag*, *Pol*, *Env* of human endogenous retrovirus subfamily H (HERV-H) (**Fig. 1a**). From dCas9 ChIP-seq, we found that LTR7 CARGO can target dCas9-TurboID to ∼1,600 copies of LTR7 loci (73.5% of the total) in the human genome (**Fig. 1c, 1d** and **Extended Data Fig. 1a**). After three hours of exogenous biotin treatment, we observed distinct biotinylation patterns in nuclear proteins of hESCs stably expressing dCas9-TurboID CARGO sgRNAs, versus hESCs expressing control sgRNAs (**Fig. 1e**). Mass spectrometry identified 144 candidate LTR7-bound proteins that were highly enriched (Fold change > 1.5, P < 0.05) among biotinylated proteins in the dCas9-TurboID/CARGO-expressing cells compared to the control cells (**Fig. 1f** and **Supplementary Table 2**). These proteins include those involved in histone modification, chromosome organization, transcription initiation, mRNA processing, and signaling related to the pluripotency factors POU5F1 and NANOG (**Fig. 1g** and **Supplementary Table 3, 4**). The enrichment of these proteins at LTR7 sequences is consistent with their known functions as active enhancers, promoters, the binding sites of pluripotency transcription factors, and boundary sequences of topologically associating domains (TAD) in hESCs^41–44^.

### The RNA m^6^A reader, YTHDC2, binds to LTR7 loci and limits transcriptional silencing of LTR7/HERV-H in a YTH-domain-dependent manner

Intriguingly, several proteins involved in the RNA methylation process were also enriched on LTR7 loci, including YTHDC2, an RNA m^6^A-reader that contains the m^6^A-binding YTH domain^22, 31^ (**Fig. 1f, 1g**). Streptavidin-pulldown followed by Western blot also indicated that YTHDC2 was highly enriched in biotinylated proteins spatially proximal to the dCas9-TurboID/CARGO ribonucleoprotein complex (**Extended Data Fig. 2a**), suggesting genomic occupancy of YTHDC2 at LTR7 loci. To directly test this idea, we conducted ChIP-seq in hESCs using an antibody directed against YTHDC2, and found that YTHDC2 was highly enriched in the LTR7 regions, that are associated with the active chromatin mark histone H3K27ac and high chromatin accessibility (ATAC-seq) (**Fig. 2a, 2b**). Sequences enriched for YTHDC2 binding also showed high chromatin accessibility, enrichment of active chromatin marks such as histone H3K27ac and DNA hypomethylation, and binding of the transcription activator, BRD4 (**Extended Data Fig. 2b**)^45, 46^. By contrast, there was minimal representation of the repressive heterochromatin mark H3K9me3 in these sequences (**Extended Data Fig. 2b**). Notably, the occupancy profile of YTHDC2 is drastically different from what has been identified for YTHDC1, another m^6^A reader that has been shown to be enriched in repressive heterochromatin loci in recent studies^22, 47, 48^. Together, our experiments indicate that LTR7 loci are enriched for YTHDC2 binding, and that genomic loci bound by YTHDC2 are enriched for active chromatin marks.

**Figure 2.**
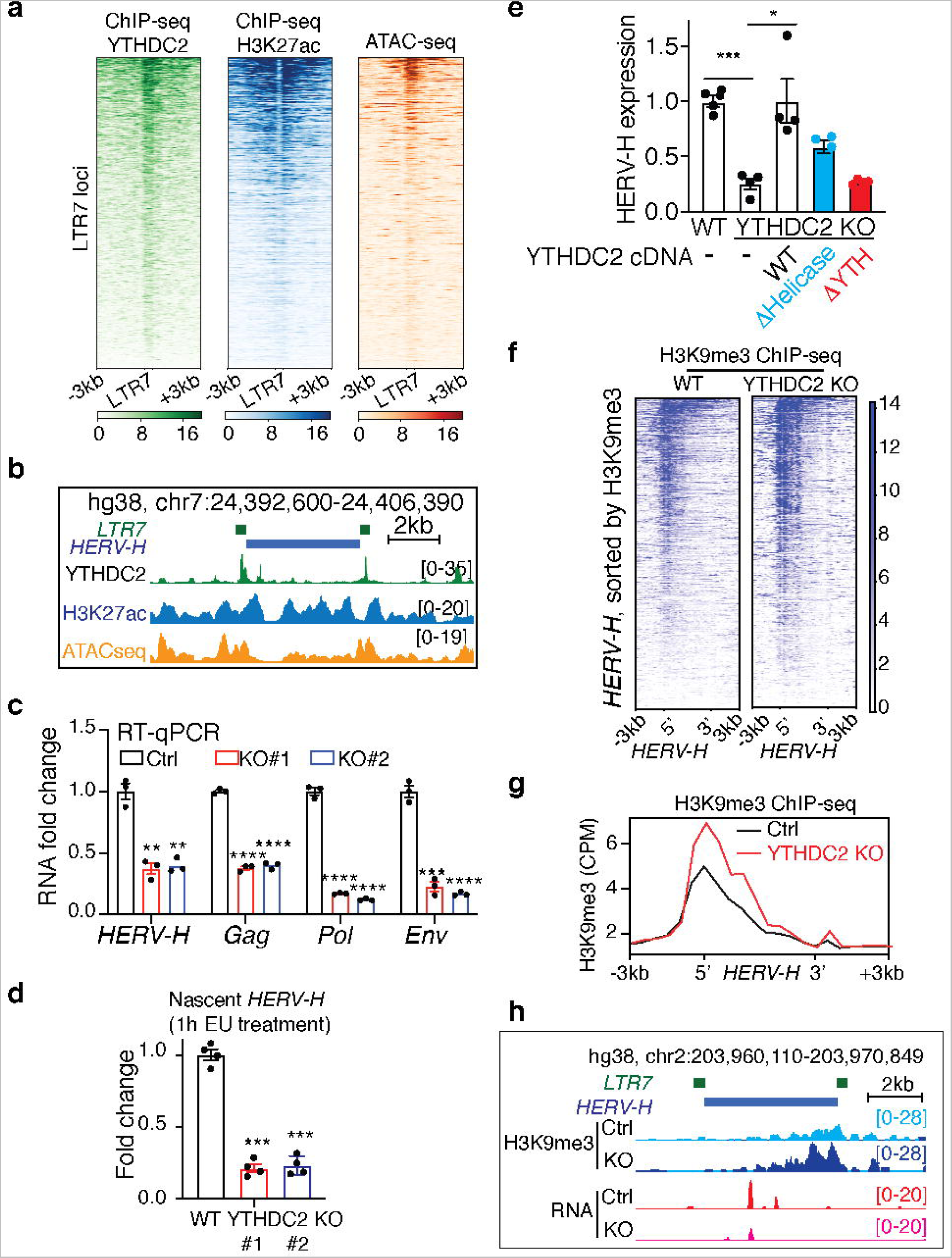
YTHDC2 occupies LTR7 genomic loci and limits their transcriptional silencing. **a, b,** Heatmaps (**a**) and UCSC genome browser snapshot (**b**) showing YTHDC2 ChIP-seq, H3K27ac ChIP-seq and ATAC-seq signal enrichment on LTR7s. **c,** RT-qPCR detection of HERV-H and viral genes expression in wild-type (WT) control versus YTHDC2 KO hESCs. **d,** RT-qPCR detection of newly transcribed HERV-H labeled with EU in WT and YTHDC2 KO cells. **e,** RT-qPCR detection of HERV-H expression level in WT hESCs, and the YTHDC2 KO hESCs expressing the indicated wild-type and mutant YTHDC2 cDNA constructs that lacks helicase activity, and m^6^A binding activity (YTH domain mutation). **f-h,** Heatmaps (**f**), aggregated chromatin signals (**g**) and UCSC genome browser snapshot (**h**) showing histone H3K9me3 ChIP-seq signal in WT and YTHDC2 KO hESCs. Data represent mean ± SEM from three independent experiments. *p < 0.05, **p < 0.01,***p < 0.001. P values were calculated by two-tailed Student’s test (**c-e**).

To interrogate YTHDC2’s function in hESCs, we generated two independent YTHDC2 knock-out (KO) hESC clones using CRISPR/Cas9 editing (**Extended Data Fig. 2c**). Interestingly, RNA levels of *HERV-H*, as well as its retroviral genes *Gag*, *Pol,* and *Env,* were significantly reduced in YTHDC2 KO clones compared to the wild-type control hESCs (**Fig. 2c**). It is well-established that *HERV-H* expression depends on the pluripotent self-renewal state of hESCs, and that m^6^A modification regulates the stability of RNA transcripts^49^. To test if the reduction of *HERV-H* expression was due to loss of pluripotency or destabilization of HERV-H transcripts, we examined the expression of the pluripotency markers *POU5F1*, *SOX2*, and *NANOG* at RNA (**Extended Data Fig. 2d**) and protein (**Extended Data Fig. 2e**) levels, and assessed the RNA decay rate of *HERV-H* RNAs (**Extended Data Fig. 2f**) in both YTHDC2 mutant and wild-type hESCs. These results suggest that the reduction of *HERV-H* RNAs in YTHDC2 KO hESCs is caused by reduced transcription. Indeed, 5-Ethynyluridine (EU) to label newly synthesized RNAs^50^ indicated that there are far fewer (∼20%) nascent HERV-H transcripts in YTHDC2 KO hESCs than in control cells (**Fig. 2d**).

Phenotypes in two independent YTHDC2 KO H1 hESC clones were similar (**Fig. 2c, 2d** and **Extended Data Fig. 2c-2f**), and we selected one representative clone for further in-depth analysis. To examine whether reduced HERV-H transcription in this mutant might be due to CRISPR/Cas9 off-target editing, we performed a rescue experiment with re-expression of wild-type YTHDC2 cDNA (**Extended Data Fig. 2g**). YTHDC2 reintroduction restored the expression of HERV-H (**Fig. 2e**), indicating limited if any effects of potential Cas9 off-target mutations. YTHDC2 was initially discovered as an RNA m^6^A reader^31^, but recent studies have shown that the protein can function as an RNA helicase in the context of spermatogenesis, independent of its m^6^A binding activity^51–53^. To determine whether the m^6^A binding activity is required for YTHDC2 to regulate HERV-H, we overexpressed in YTHDC2 KO hESCs two mutant forms of YTHDC2 that lack either helicase or m^6^A binding activity (**Fig. 2e**, helicase Δ with ATP-binding site mutation, and ΔYTH with YTH-domain deletion, respectively. **Extended Δ Data Fig. 2g**). Interestingly, the YTHDC2^Δhelicase^ mutant could partially rescue HERV-H expression in YTHDC2 KO cells, whereas the YTHDC2 ^YTH^ mutant had no detectable rescue activity. These data indicate that the m^6^A binding capacity of YTHDC2 is indispensable for its regulation of HERV-H transcription, and that its helicase activity also contributes.

To determine global genomic features associated with YTHDC2 depletion in hESCs, we assessed transcriptome and chromatin signatures of YTHDC2 KO hESCs. Consistent with our observation from RT-qPCR analysis (**Fig. 2c**), RNA-seq results also revealed significantly decreased HERV-H expression in the absence of YTHDC2 in hESCs (**Extended Data Fig. 2h**, Wilcoxon *P* < 2.5e-16). In eukaryotic cells, the inactivation of TE chromatin is associated with the deposition of repressive heterochromatic histone marks such as H3K9me3 ^5^. Indeed, results from H3K9me3 ChIP-seq data showed a significant gain of histone H3K9me3 modification (**Fig. 2f-2h**) on LTR7/HERV-H loci in YTHDC2 KO hESCs compared to wild-type cells. These results demonstrate that YTHDC2 is required to prevent epigenetic silencing of LTR7/HERV-H chromatin loci.

### The YTHDC2/LTR7 axis causally regulates the switch between pluripotency versus the neuronal fate of hESC

Previous studies have reported that RNAi-mediated knockdown of HERV-H RNAs causes loss of pluripotency and spontaneous differentiation of hESCs^54, 55^. However, despite a significant reduction of HERV-H RNAs in the YTHDC2 KO hESCs, the mutant hESCs didn’t lose their pluripotency in the self-renewal medium (**Extended Data Fig. 2d, 2e**, considered further in the discussion section). In human iPSCs, the ectopic activation of LTR7 is associated with defects in neuronal differentiation^56, 57^. Indeed, the differentially expressed genes of hESCs in the absence of YTHDC2 are enriched for Gene Ontology terms related to nervous system development (**Extended Data Fig. 3a**). From YTHDC2 ChIP-seq, we found that the binding sites of YTHDC2 were also enriched in the sequences nearby the genes related to neurogenesis and nervous system development (**Extended Data Fig. 3b**). These results suggest the potential role of YTHDC2 in regulating neuronal fate.

**Figure 3.**
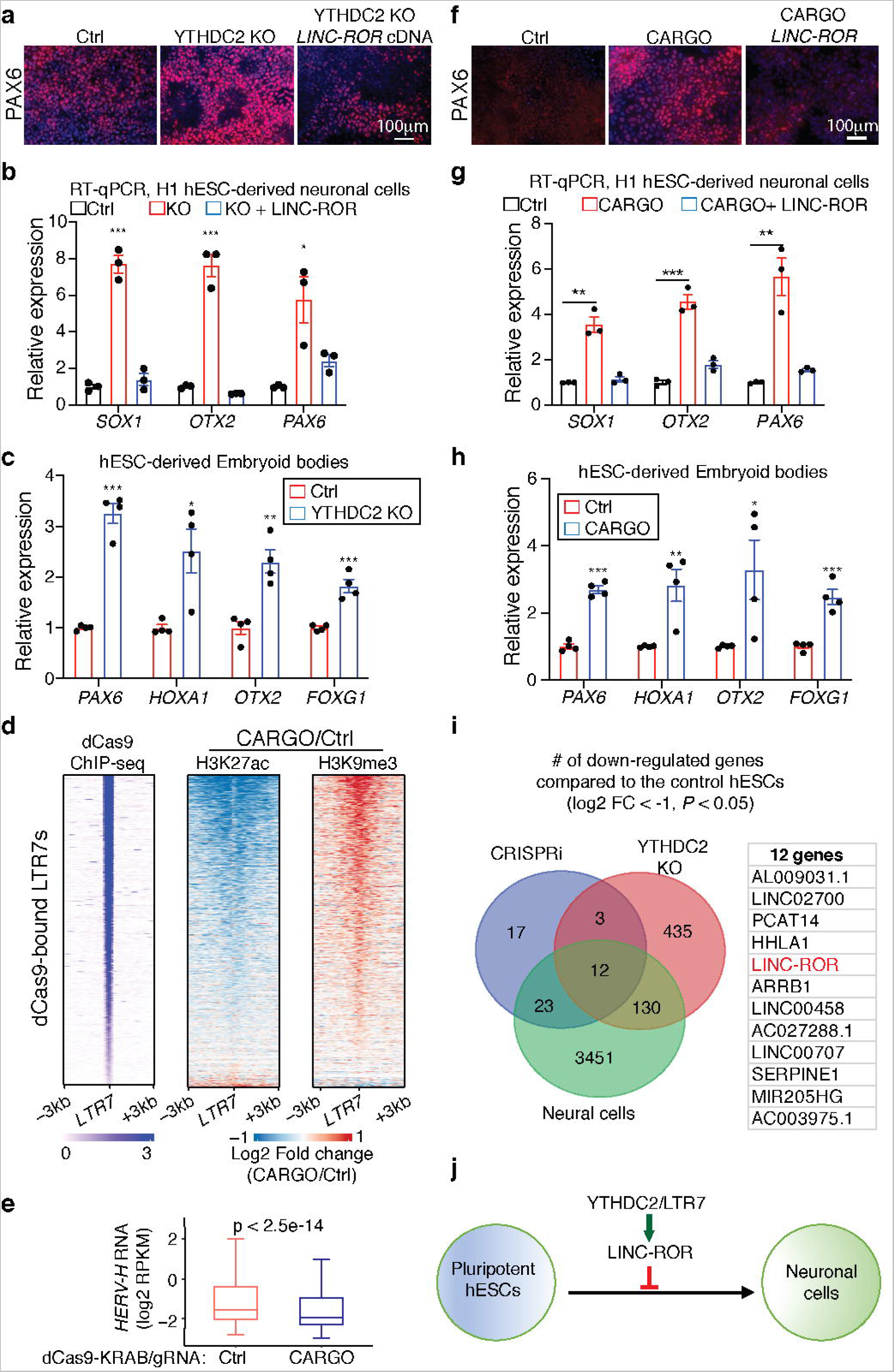
The YTHDC2/LTR7 axis inhibits neuronal fate of hESCs via *LINC-ROR*. **a, b,** The hESCs that are wild-type, YTHDC2 KO, and YTHDC2 KO with LINC-ROR overexpression were cultured in the neural induction media to induce lineage specific neuronal differentiation. (**a**) Immunofluorescence staining of neuronal cell marker PAX6, and (**b**) RT-qPCR analysis of the indicated neuronal cell markers SOX1, OTX2 and PAX6. **c,** Embryoid body (EB) derived from both Ctrl and YTHDC2 KO hESCs were collected for RNA extraction, followed by RT-qPCR analysis of indicated neuronal lineage marker genes. **d,** dCas9-KRAB and the LTR7 targeting CARGO construct were co-expressed in hESCs to induce epigenetic silencing of LTR7 genomic loci. Heatmaps showing log2 fold change of H3K27ac and H3K9me3 ChIP-seq signals in LTR7-silenced hESCs versus control hESCs expressing dCas9-KRAB and non-targeting control sgRNA. Each line represents a LTR7 locus (+/-3kb of LTR7 annotation) that is bound by dCas9. **e,** Boxplots showing HERV-H expression level detected by RNA-seq in Ctrl and LTR7-silenced hESCs. **f, g,** The hESCs that are wild-type, LTR-silenced, and LTR7-silenced with LINC-ROR overexpression, were cultured in the neural induction media to induce lineage specific neuronal differentiation. (**f**) Immunofluorescence staining of neuronal cell marker PAX6, and (**g**) RT-qPCR analysis of the indicated neuronal cell markers SOX1, OTX2, and PAX6. **h**, Embryoid body (EB) derived from both control and LTR7-silenced hESCs were collected for RNA extraction, and RT-qPCR analysis of indicated neuronal lineage markers. **i**, (Left) Venn diagrams showing the overlap of downregulated genes in LTR7-silenced hESCs, YTHDC2 KO, and hESC-derived neuronal cells compared to the wild-type control hESCs cultured in the self-renewal media (log_2_FC > 1, *P* < 0.05). (Right) The list of 12 RNA transcripts that are down-regulated in the three conditions. The HERV-H derived LINC-ROR is highlighted in red. **j**. Schematic illustration showing the YTHDC2/LTR7-axis prevents neural fate of pluripotent hESCs by regulating LINC-ROR expression. Data represent mean ± SEM from three independent experiments. *p < 0.05,** < 0.01, ***p < 0.001. P values were calculated by two-tailed Student’s test (**b, c, g** and **h**) and Wilcoxon signed-rank test (**e**).

To determine the impact of YTHDC2/LTR7 depletion on neurogenic potential of hESCs, we used two complementary systems: a lineage-specific neuronal cell differentiation assay, based on neural induction medium that contains SMAD2/3 inhibitors^58^, and an embryoid-bodies (EBs) formation assay that allows spontaneous differentiation of hESC towards multiple lineages. In the neural-induction medium, we found that YTHDC2 KO hESCs express ∼6-8 fold higher levels of neuronal marker genes than wild-type cells, as indicated by immunostaining of PAX6 (**Fig. 3a**) and RT-qPCR analysis of *SOX1, OTX2*, and *PAX6* transcripts (**Fig. 3b**). Additionally, embryoid-bodies (EBs) derived from YTHDC2 KO cells also displayed significantly higher levels (∼2-3 folds) of neuronal markers such as *PAX6*, *HOXA1*, *OTX2*, and *FOXG1* (**Fig. 3c**). These data indicate that YTHDC2 presence controls the ability of pluripotent stem cells to differentiate toward the neuronal lineage.

Next, to determine whether LTR7 sequences themselves influence the fate of hESCs, we induced epigenetic silencing of more than ∼1,600 copies of LTR7 sequences *en mass* by co-expression of the epigenetic repressor dCas9-KRAB^59^ together with LTR7 CARGO in hESCs^32, 33^. ChIP-seq revealed that dCas9-KRAB/CARGO-targeted LTR7 sequences (73.5% of the total LTR7s) exhibited global loss of active chromatin mark H3K27ac and gain of repressive mark H3K9me3 (**Fig. 3d** and **Extended Data Fig. 3c** for genome browser visualization), indicating effective epigenetic perturbation of LTR7 loci. Concomitantly, this perturbation of LTR7 loci caused a significant reduction of *HERV-H* transcripts (**Fig. 3e**, RNA-seq, Wilcoxon *P* < 2.5e-14; **Extended Data Fig. 3d** RT-qPCR;). Intriguingly, epigenetic silencing of LTR7 loci did not cause loss of pluripotency or induce hESC differentiation, a phenotype that was reported in the previous RNAi-mediated HERV-H knock-down studies (considered further in discussion)^54, 55^. Instead, key observations indicate that LTR7 silencing phenocopied YTHDC2 KO in hESCs. First, LTR7-silenced hESCs were maintained in a pluripotent state in self-renewal medium, as determined by cell morphology (**Extended Data Fig. 3e**) and the expression of pluripotency markers, including *POU5F1, SOX2* and *NANOG* at both the RNA (**Extended Data Fig. 3f**, RT-qPCR) and protein (**Extended Data Fig. 3g**, immunostaining) levels. Second, they expressed a higher level (∼3-6 fold) of neuronal makers in both lineage-specific neural-induction medium (**Fig. 3f** immunostaining of PAX6; **Fig. 3g**, RT-qPCR of *PAX6, SOX1, OTX2*) and in the EB formation assay (**Fig. 3h**, RT-qPCR, *PAX6, HOXA1, OTX2, FOXG1*). Taken together, our results suggest a model in which the YTHDC2/LTR7 axis is dispensable for pluripotency maintenance but negatively regulates the switch from pluripotency to the neuronal fate.

To identify key regulators downstream of YTHDC2/LTR7 responsible for the fate switch of hESCs, we examined genes with significantly lowered levels in each condition of YTHDC2 KO hESC, LTR7-silenced hESCs, and hESC-derived neuronal cells, as compared to control pluripotent hESCs (**Fig. 3i**, RNA-seq DESeq2 log 2 (fold change) > 1, *P* value < 0.05, **Supplementary Table 5-7**). This comparison identified 12 genes and long non-coding RNAs, including *LINC-ROR,* one of the LTR7/HERV-H-derived long non-coding RNAs that was initially identified as a key regulator in human iPSC reprogramming^60^. *LINC-ROR* can maintain hESC self-renewal by acting as a microRNA sponge to trap miR-145, which would otherwise target multiple core pluripotency factors including *POU5F1*, *NANOG*, and *SOX2*^61, 62^. RT-qPCR assessment indicated that *LINC-ROR* levels were indeed significantly reduced in YTHDC2 KO hESCs and in LTR7-silenced hESCs (**Extended Data Fig. 3h**). To test if increases in *LINC-ROR* could impact the bias towards neuronal differentiation caused by knockout of YTHDC2 or silencing of LTR7, we overexpressed the cDNA in hESCs. We found that this manipulation reduced the expression of neuronal markers in hESCs when cultured in the neural induction medium (**Fig. 3a, 3f**, PAX6 immunostaining on the right panel; **Fig. 3b, 3g**, RT-qPCR of *SOX1*, *OTX2*, and *PAX6*). Considering previous studies implicating *LINC-ROR* in iPSC reprogramming and pluripotency maintenance, our data suggest that the YTHDC2/LTR7 axis reinforces pluripotency and limits neural differentiation of hESCs, at least partially through control of *LINC-ROR* (**Fig. 3j**).

### m^6^A-modified HERV-H RNAs recruit YTHDC2 to LTR7/HERV-H genomic loci

What is the underlying mechanism that directs the m^6^A reader YTHDC2 to LTR7 genomic loci? As we had observed that the m^6^A binding, YTH-domain of YTHDC2 is required for its regulation of LTR7/HERV-H, we hypothesized that YTHDC2 interacts with m^6^A-modified RNAs in the vicinity of LTR7 DNA (**Fig. 4a**). *HERV-H* transcripts are expressed from their LTR7 promoters, supporting the feasibility of this mechanism. To test this idea, we conducted RNA immunoprecipitation sequencing (RIP-seq) analysis to integrate the m^6^A modification and YTHDC2 binding of HERV-H transcripts. From published hESC m^6^A RIP-seq^49^ and our own hESC YTDHC2 RIP-seq data, we found that HERV-H transcripts were indeed extensively modified by m^6^A (**Fig. 4b**) and bound by YTHDC2 (**Fig. 4c**, and **Extended Data Fig. 4a**). Notably, the RNA regions bound by YTHDC2 were enriched for the m^6^A consensus motif GGAC (RRACH)^31^ (**Extended Data Fig. 4b**), suggesting YTHDC2 function as an m^6^A reader in hESCs. To further interrogate the physical interaction between HERV-H RNA and LTR7 DNA, we re-analyzed published hESC MARGI (MApping RNA-Genome Interactions) data, which enables quantitative detection of genome-wide RNA-chromatin interactions via proximity ligation of chromatin-associated RNAs with their target genomic sequences^63^. This analysis revealed that HERV-H RNAs form extensive and significant interactions with LTR7 DNA sequences (**Fig. 4d**). Together with YTHDC2 ChIP-seq data, these results strongly support the notion that physical interactions occur between YTHDC2, m^6^A-modified HERV-H RNAs, and LTR7 DNA sequences in hESCs.

**Figure 4.**
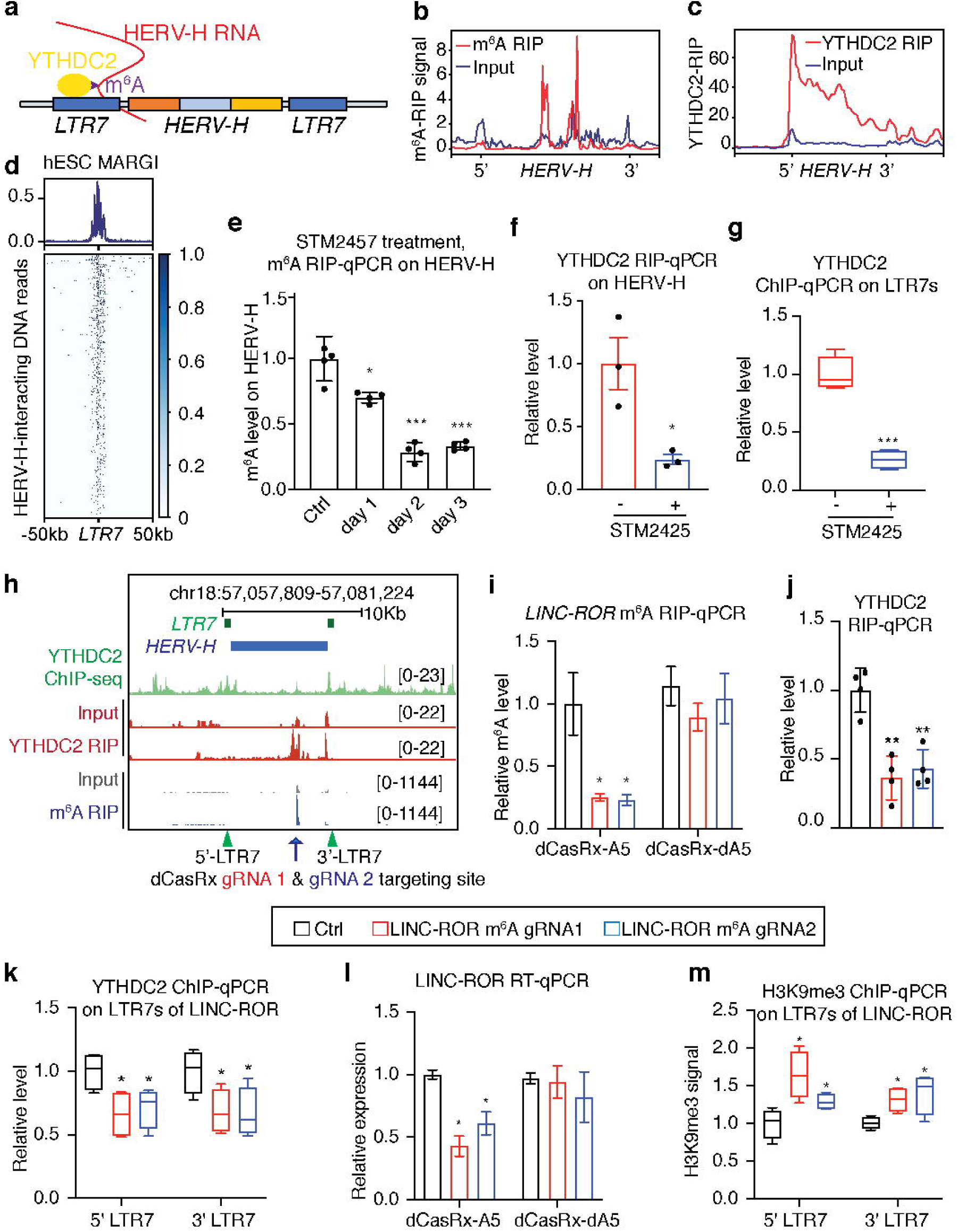
The m^6^A-modified HERV-H RNAs recruit YTHDC2 to LTR7 genomic loci in a m^6^A dependent manner. **a,** Schematic illustration of the proposed model: YTHDC2 is recruited to the LTR7/HERV-H DNA sequences via HERV-H RNAs that are m^6^A modified and interact with LTR7 DNA regions. **b, c,** Aggregated RIP-seq signal of m^6^A MeRIP (**b**) and YTHDC2 RIP-seq (**c**) over the HERV-H RNAs. **d,** Heatmap (bottom) and the aggregated reads counts (top) showing that the HERV-H RNAs bind to LTR7 genomic loci. Each row of the heatmap represents one LTR7 locus. **e,** m^6^A MeRIP-qPCR quantification of the m^6^A levels of HERV-H RNAs in DMSO and STM2457 treated hESCs at various time points. **f, g,** YTHDC2 RIP-qPCR (**f**) and ChIP-qPCR (**g**) quantification of YTHDC2 binding on HERV-H transcripts and LTR7 genomic loci, respectively, in hESCs after 2 days of DMSO (control) and STM2457 treatment. **h,** UCSC genome browser snapshot of LINC-ROR locus, showing YTHDC2 ChIP-seq, YTHDC2 RIP-seq, and m^6^A MeRIP-seq tracks. Two gRNAs were designed to target this m^6^A peak. **i,** m^6^A MeRIP-qPCR showing reduced m^6^A level of LINC-ROR by dCasRx-ALKBH5 but not by dCasRx-dALKBH5. **j, k,** YTHDC2 RIP-qPCR (**j**) and ChIP-qPCR (**k**) quantification of YTHDC2 binding on HERV-H transcripts and LTR7 genomic loci, respectively, upon m^6^A removal. **l,** RT-qPCR detection of LINC-ROR expression level in hESC expressing dCasRx-ALKBH5 versus dCasRx-dALKBH5 with Ctrl or LINC-ROR targeting gRNAs. **m**, H3K9me3 ChIP-qPCR analysis showing the changes of H3K9me3 signal on the two LTR7 loci of LINC-ROR after m^6^A editing on LINC-ROR. dCasRx-A5: dCasRx-ALKBH5; dCasRx-dA5: dCasRx-dALKBH5. Black bars: non-targeting control gRNA; red and blue: the two guide RNAs (#1 and #2) targeting the m6A-modified sequence of LINC-ROR (**i-m**). Data represent mean ± SEM from three independent experiments. *p < 0.05, ** p < 0.01, ***p < 0.001. P values were calculated by two-tailed Student’s test.

Next, we asked whether the m^6^A modification of HERV-H is required for the binding of YTHDC2 on LTR7/HERV-H transcripts and DNA sequences. We first reduced global m^6^A modification of hESCs by treating them with STM2457^64^, a highly potent and selective catalytic inhibitor of the m^6^A writer METTL3. Two days after STM2457 treatment, the m^6^A level of all HERV-H RNAs was decreased by ∼80% (**Fig. 4e**). Notably, we found that STM2457 treatment did not induce HERV-H RNA degradation (**Extended Data Fig. 4c**), but disrupted the binding of YTHDC2 on both HERV-H transcripts (**Fig. 4f**) and LTR7 genomic loci (**Fig. 4g**).

To address the potential concern that loss of YTHDC2 binding at LTR7/HERV-H loci is due to the indirect effects of global alterations of transcriptome and epitranscriptome by STM2457 treatment, we next conducted a sequence-specific m^6^A editing experiment using *LINC-ROR* as a representative locus. This was selected because (1) *LINC-ROR* RNA is modified by m^6^A and bound by YTHDC2; and (2) YTHDC2 bind to DNA sequences of the two LTR7s flanking the HERV-H sequence of LINC-ROR (**Fig. 4h**). To edit m^6^A modifications of *LINC-ROR*, we fused the m^6^A-eraser ALKBH5 with dCasRx (dead CasRx, the RNA-guided RNA-targeting type VI CRISPR-Cas^65, 66^) to construct dCasRx-ALKBH5, which can remove m^6^A from its target RNA in a gRNA-dependent manner^67, 68^. We designed two gRNAs targeting the m^6^A-modified sequence of *LINC-ROR* (**Fig. 4h**, blue arrow), to guide dCasRx-ALKBH5 and its mutant form dCasRx-dALKBH5 (dead ALKBH5) lacking m^6^A-eraser activity (**Fig. 4i**, dCasRx-A5 versus dCasRx-dA5), to the m^6^A-modified sequence of *LINC-ROR*. We found that dCasRx-ALKBH5, but not dCasRx-dALKBH5, can effectively reduce the extent of m^6^A modification of *LINC-ROR* (**Fig. 4i**). Significantly, the removal of m^6^A from *LINC-ROR* resulted in a markedly decrease of YTHDC2 binding on *LINC-ROR* RNA (**Fig. 4j**, RIP-qPCR) and on the two LTR7 loci of *LINC-ROR* (**Fig. 4k**, ChIP-qPCR). Taken together, we conclude that the m^6^A modification of HERV-H transcripts is required for the binding of YTDHC2 on both HERV-H RNAs and LTR7/HERV-H DNA sequences.

### The m^6^A modification of HERV-H RNAs is required to prevent LTR7/HERV-H silencing in a YTHDC2-dependent manner

Next, we aimed to determine whether m^6^A modification of HERV-H transcripts is required to prevent the silencing of LTR7/HERV-H loci. Using *LINC-ROR* locus as a paradigm, we found that the removal of its m^6^A led to a significant reduction of *LINC-ROR* RNA (**Fig. 4l**), which was not due to the destabilization of *LINC-ROR* transcripts (**Extended Data Fig. 4d**) or the binding of dCasRx-dALKBH5 (**Fig. 4l**). Furthermore, we found that upon removal of m^6^A from *LINC-ROR* RNA, the genomic locus of *LINC-ROR* exhibited a significant gain of the repressive heterochromatin marks histone H3K9me3 (**Fig. 4m**), indicating epigenetic silencing. In YTHDC2 KO hESCs, we conducted the same m^6^A editing experiment and found that in the absence of YTHDC2, the removal of m^6^A from *LINC-ROR* did not induce a further reduction of *LINC-ROR* or gain of H3K9me3 modification on its LTR7 loci (**Extended Data Fig. 4e-4g**). These data indicate that the m^6^A modification of *LINC-ROR* regulates its own transcription in a YTDHC2-dependent manner. Collectively, our data support a model in which the m^6^A modification of HERV-H RNAs plays a causal role in preventing the epigenetic silencing of LTR7/HERV-H genomic loci by recruiting the RNA m^6^A reader YTHDC2 (**Fig. 4a**).

### YTHDC2 recruits TET1 for DNA demethylation of LTR7/HERV-H genomic loci

Having established a causal role of m^6^A-modified HERV-H RNAs in preventing LTR7/HERV-H silencing by recruiting YTHDC2, we further interrogated the mechanisms by which YTHDC2 is required for LTR7/HERV-H activation. From CARGO BioID data, both TET1 and TET3 were identified as candidate proteins enriched on LTR7 loci in hESCs (**Fig. 1g**). In mammals, TET enzymes oxidize 5-methylcytosine (5mC) to 5-hydroxymethylcytosine (5hmC), which leads to progressive loss of DNA methylation and activation of several TE families and *cis*-regulatory sequences^69–73^. A previous study showed that genetic depletion of TET1, but not TET2 or TET3, leads to a marked decrease of global 5hmC by ∼80% in hESCs^74^, suggesting that TET1 plays a primary role to control 5hmC-mediated DNA demethylation in hESC. Thus, we asked whether TET1 proteins play a role in the YTDHC2-mediated regulation of LTR7/HERV-H.

Using a CRISPR/Cas9-mediated knock-in strategy described recently^75^, we fused a V5 epitope to the N-terminus of the endogenous TET1 protein, in hESC clones with intact or disrupted YTHDC2 genes, to enable specific immunoprecipitation of endogenous TET1. This enabled us to investigate the protein-protein interactions and genomic occupancy of TET1 and YTHDC2 in hESCs. Based on endogenous protein co-immunoprecipitation experiments, we determined that YTHDC2 and TET1 proteins interact with each other in hESC (**Fig. 5a**). By comparing TET1-V5 and YTHDC2 ChIP-seq data, we found that TET1 was significantly enriched on the LTR7s loci occupied by YTHDC2, and vice versa (**Fig. 5b**). Genome-wide quantification further revealed that the ChIP-seq signals of TET1 and YTHDC2 were highly correlated with each other across all the LTR7 loci (**Extended Data Fig. 5a**, Pearson correlation coefficient = 0.61, *P* < 0.001). These results indicate that TET1 and YTHDC2 interact with each other and also co-localize at LTR7 loci, suggesting their cooperative regulation of LTR7 activity in hESCs. Importantly, the binding of TET1 on LTR7 sequences was markedly decreased in YTHDC2 KO cells (**Fig. 5c**), accompanied by a significant gain of DNA 5mC modification and a marked decrease of 5hmC modification surrounding the LTR7/HERV-H loci (**Fig. 5d, 5e** and **Fig. 5f** for genome browser visualization of the results shown in **Fig. 5b-5e**). These data strongly suggest that TET1 is recruited by YTHDC2 to the LTR7/HERV-H loci where it induces DNA CpG demethylation of TE sequences.

**Figure 5.**
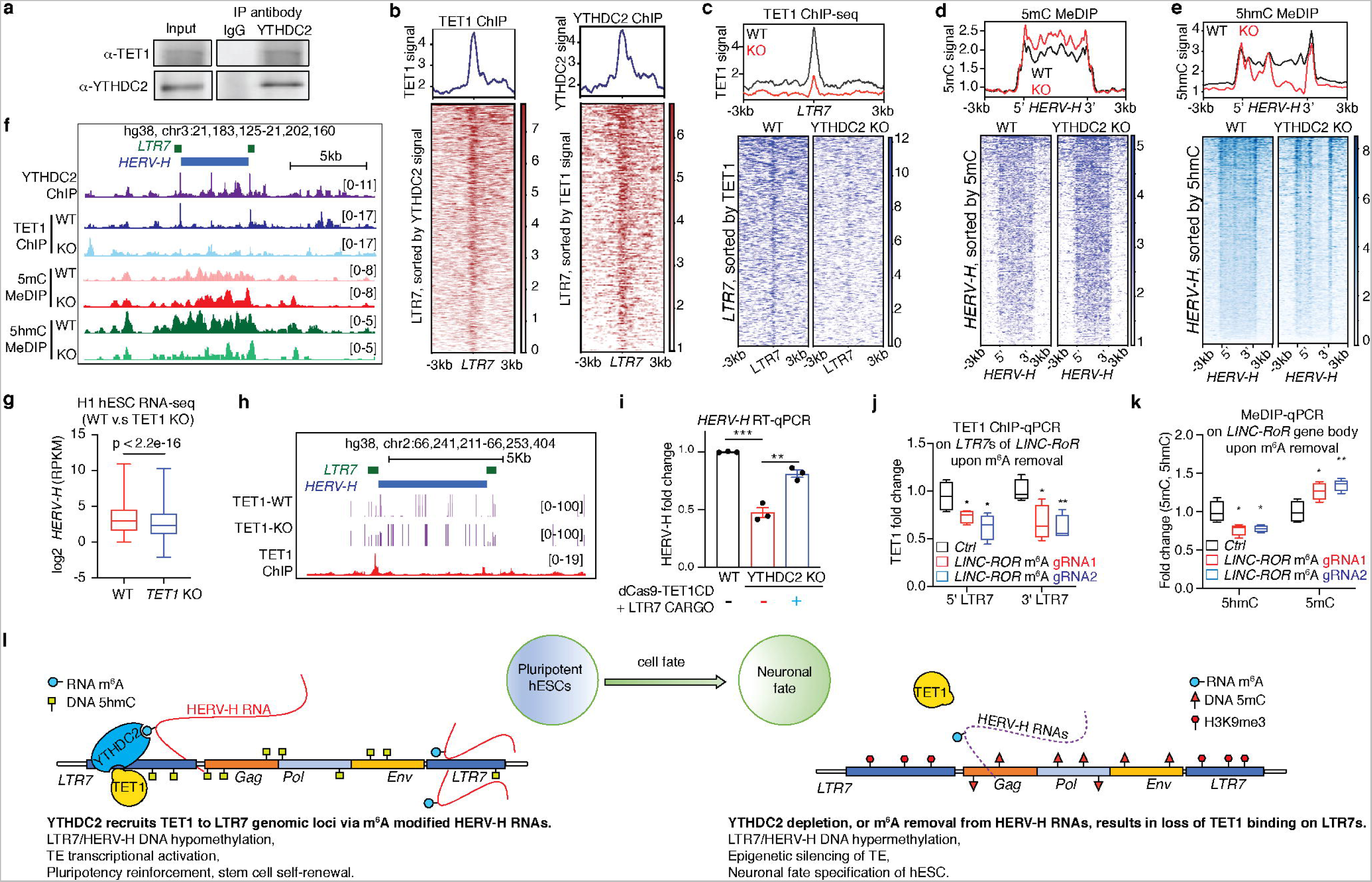
YTHDC2 recruits TET1 to prevent epigenetic silencing of LTR7/HERV-H loci via DNA demethylation. **a,** Endogenous protein co-immunoprecipitation showing YTHDC2 interacts with TET1 in H1 hESCs. Representative results from three independent experiments are shown. **b,** Heatmap showing the endogenous TET1-V5 ChIP-seq signal centered on the LTR7 loci that are bound by YTHDC2 (left) and showing YTHDC2 ChIP-seq signal centered on the LTR loci that are bound by TET1-V5 (right). **c,** Heatmaps (bottom) and reads counts coverage (top) of TET1 ChIP-seq on the LTR7 loci in wild-type (WT) versus YTHDC2 KO hESCs (KO). The LTR7 loci are sorted based on their TET1 binding strength in the WT hESCs. **d,** Heatmaps (bottom) and reads counts coverage (top) of 5mC MeDIP on the LTR7/HERV-H loci in wild-type (WT) versus YTHDC2 KO hESCs (KO). The LTR7/HERV-H loci are sorted based on their DNA 5mC signal in the YTHDC2 KO hESCs. **e,** Heatmaps (bottom) and reads counts coverage (top) of 5hmC MeDIP on the LTR7/HERV-H loci in wild-type (WT) versus YTHDC2 KO hESCs (KO). The LTR7/HERV-H loci are sorted based on their DNA 5hmC signal in the WT hESCs. **f,** UCSC genome browser snapshot showing a representative LTR7/HERV-H locus exhibiting loss of TET1 binding (ChIP-seq), gain of 5mC (MeDIP-seq) signal and loss of 5hmC (MeDIP-seq) signal on LTR7 loci in WT versus YTHDC2 KO hESCs. **g,** RNA-seq data showing the reduction of HERV-H RNAs in TET1 KO H1 hESCs compared to the control wild-type H1 hESCs. **h,** UCSC genome browser snapshot showing a representative LTR7/HERV-H locus exhibiting gain of methylation upon TET1 knockout. **i,** HERV-H levels are quantified by RT-qPCR in hESCs that are WT, YTHDC2 KO, and YTHDC2 KO co-expressing dCas9-TET1CD with LTR7 CARGO. TET1CD: TET1 catalytic domain. **j,** TET1 ChIP-qPCR analysis showing the changes of TET1 signal on the two LTR7 loci of LINC-ROR after m^6^A editing on LINC-ROR. **k,** DNA 5hmC and 5mC MeDIP-qPCR analysis showing the changes of 5hmC and 5mC signals on the gene body of LINC-ROR after m^6^A editing on LINC-ROR. **l,** Cartoon illustration of the model proposed in this study: YTHDC2 recruits TET1 to the genomic loci of LTR7/HERV-H, through its interaction with m^6^A modified HERV-H RNAs. The engagement of TET1 on LTR7 leads to DNA demethylation of TE loci, which prevents epigenetic silencing of LTR7/HERV-H chromatin and inhibits the neuronal fate of hESCs. Data represent mean ± SEM from three independent experiments. * p < 0.05, ** p < 0.01, ***p < 0.001. P values were calculated by two-tailed Student’s test (**i-k**) and Wilcoxon signed-rank test (**g**).

Next, to determine whether TET1 causally regulates the activity of LTR7/HERV-H, we analyzed the published RNA-seq and methyl capture sequencing data generated from wild-type and TET1 KO H1 hESCs^75^. We found that TET1 KO indeed led to a significant reduction of HERV-H expression in H1 hESC (**Fig. 5g**, Wilcoxon p < 2.2e-16), accompanied by a gain of DNA CpG methylation in LTR/HERV-H sequences (Fig. 5h). Notably, the reduced HERV-H expression in the YTHDC2 KO hESCs can be rescued by overexpression of LTR7 CARGO together with TET1CD, a fusion protein containing dCas9 and the catalytic domain (CD) of TET1 that can convert DNA 5mC to 5hmC on its targeted chromatin regions^76^ (**Fig. 5i**). Collectively, these data establish the causal role of TET1 in controlling LTR7/HERV-H activity in hESCs, and show that the silencing of LTR7/HERV-H in the absence of YTHDC2 is due at least in part to the loss of TET1 binding on LTR7/HERV-H DNA.

### The m^6^A modification of HERV-H is required for TET1 binding and LTR7/HERV-H DNA demethylation

Lastly, we asked whether m^6^A modification of HERV-H RNA is required for TET1 engagement on LTR7/HERV-H DNA and conversion of 5mC to 5hmC. We found that upon targeted removal of m^6^A from *LINC-ROR* RNA (**Fig. 4h, 4i**), the binding of TET1 on the two LTR7 loci of *LINC-ROR* was markedly decreased (**Fig. 5j**) the hESCs with intact YTHDC2 expression. By contrast, erasing m^6^A from *LINC-ROR* in the absence of YTHDC2 did not lead to further reduction of TET1 binding on the *LINC-ROR* locus (**Extended Data Fig. 5b**). These data strongly suggest that the recruitment of TET1 to the LTR7/HERV-H DNA requires both m^6^A modification of HERV-H RNA and YTHDC2. Importantly, the loss of TET1 binding on the LINC-ROR locus is accompanied by a concordant gain of DNA 5mC and loss of DNA 5hmC across *LINC-ROR* sequences (**Fig. 5k** and **Extended Data Fig.5c, 5d**). These data further support a model in which m^6^A modification of HERV-H transcripts recruits TET1 to prevent epigenetic silencing of LTR7/HERV-H loci, via DNA demethylation mechanism.

In summary, through CARGO-BioID and functional analysis, we identify the RNA m^6^A reader YTHDC2 as a key regulator to prevent the silencing of LTR7/HERV-H in hESCs. YTHDC2 occupies LTR7/HERV-H loci through its interaction with m^6^A modified HERV-H transcript, and further recruits TET1 to demethylate LTR7/HERV-H DNA (**Fig. 5l**). This YTHDC2/LTR7-axis reinforces pluripotency and inhibits the neural fate of hESCs.

## DISCUSSION

In eukaryotes, RNA m^6^A modification is the most abundant epitranscriptome modification, regulating RNA splicing, stability, spatial localization, and translation ^22^. In addition, the DNA cytosine 5mC and 5hmC marks are widely distributed through the genome, determining the activity of cis-regulatory sequences that are critical for spatial-temporal gene regulation ^77, 78^. To the best of our knowledge, our work is the first study that links RNA m^6^A with DNA 5mC/5hmC regulation by demonstrating the physical and functional interaction between the RNA m^6^A reader YTHDC2 and the DNA demethylase, TET1. Despite the recent debates about whether YTHDC2 acts as an m^6^A reader ^51–53^, our data clearly show that both m^6^A modification of HERV-H RNA and the m^6^A-binding, YTH-domain of YTHDC2 are required for YTHDC2’s regulation of LTR7/HERV-H transcription and epigenetic activity. Therefore, our findings significantly expand our knowledge of the crosstalk between the epitranscriptome and the epigenome in genome regulation. Furthermore, we postulate that a similar mechanism also regulates the chromatin state of regulatory DNA sequences that are not related to TEs. Indeed, from transcriptome and epigenome data of control and YTHDC2 KO hESCs, we found that YTHDC2 and TET1 co-occupy 30,009 ChIP-seq peaks (**Extended Data Fig. 6a-6c**), representing promoters, enhancers, and other regulatory sequences of the human genome (**Extended Data Fig. 6d**). Notably, YTHDC2 depletion leads to reduced binding of TET1 and gain of H3K9me3/5mC on these shared ChIP-seq peaks (**Extended Data Fig. 6e-6j**). These results suggest that in addition to LTR7/HERV-H loci, YTHDC2 also regulates the genomic occupancy of TET1 and the chromatin state of cis-regulatory sequences in the human genome, possibly through the m^6^A modified chromatin-associated regulatory RNA (carRNAs) and mRNA of human genes (**Extended Data Fig. 6k**). Given the broad impact of m^6^A and DNA methylation on gene regulation, we expect the connection between the epitranscriptome and the epigenome discovered in this study will help our understanding of gene regulation in developmental disorders and human diseases that involve RNA m^6^A and DNA methylation.

In addition to YTHDC2 and TET1, there are another 142 proteins putatively enriched on LTR7 genomic loci, where they likely regulate LTR7/HERV-H transcription and chromatin states. In future studies, it would be interesting to determine whether and how these candidate LTR7-bound proteins collectively contribute to the robust regulation of LTR7/HERV-H transcription and chromatin epigenetic activity in pluripotency and differentiation. Notably, due to the large copy number and the highly repetitive nature of TE loci, as well as their locus-specific mutations or indels that have accumulated through evolution, it has been technically challenging to identify those effectors controlling the activity of endogenous TE sequences. Our study demonstrates that CARGO-BioID is a powerful technology to identify key regulators of TE activity. Considering the critical roles of TEs in regulating development, immunity, aging, and cancer ^1–4^, we expect that CARGO-BioID will become a powerful technology to further our understanding of TE function and regulation in many biological processes and human diseases.

One unexpected finding from this study is that the significant decrease of HERV-H transcription does not result in loss of pluripotency in YTHDC2 KO and LTR7-silenced hESCs. This result seems contradictory to previous findings showing that RNAi knock-down of HERV-H RNAs leads to the loss of pluripotency in hESC ^55^. There are two possible reasons for this discrepancy. First, many LTR7-distal human genes, located within ∼200kb of their nearest LTR7 loci, are significantly down-regulated upon LTR7 perturbation (**Extended Data Fig. 7a, 7b**). This observation is consistent with the notion that the LTR7 can act as hESC enhancers to control the expression of distal human genes^43, 79^. By contrast, these human genes regulated by distal LTR7s are less likely to be affected by shRNA-mediated HERV-H knock-down. In this scenario, it is likely that the alteration of these human genes results in a distinct phenotype compared to that of knockdown of HERV-H RNA. Second, the discrepancy could also be due to shRNA off-target effects. For example, it has been reported that the knockdown of one specific HERV-H transcript, ESRG, leads to loss of pluripotency in hESC^55^. However, in a recent study from Shiyan Yamanaka’s group ^80^, the authors showed that genetic depletion of the DNA sequence of the entire ESRG locus is dispensable for pluripotency or iPSC reprogramming ^80^. These data suggest that HERV-H knockdown causes unknown off-target effects, which in turn results in hESC differentiation and loss of pluripotency.

In conclusion, by exploiting CARGO-BioID, a TE-centric proteomic approach that we developed, we unveil previous unknown crosstalk between RNA m^6^A-modification and DNA methylation regulation and show that this under-appreciated mechanism control TE activity and cell fate switch of hESCs. Considering the significant impact of RNA and DNA methylation in regulating numerous pathobiological processes, we expect the connection between epitranscriptome and epigenome discovered in this study will throw light on genome regulation in developmental disorders and human diseases that involve RNA m^6^A and DNA methylation.

## ACCESSION CODES AND DATA AVAILABILITY

Sequencing data have been deposited to the NCBI Gene Expression Omnibus (GEO) (http://www.ncbi.nlm.nih.gov/geo) under accession number GSE210867. Other publicly available datasets used in this study include: RNA-seq of WT and TET1 knockout H1 cells (GSM5183601, GSM5183602, GSM5183607, GSM5183608), methyl capture sequencing (GSM5183585, GSM5183586, GSM5183589, GSM5183590), H3K27ac (ENCFF103PND), H3K9me3 (ENCFF385ZBQ), H3K4me1 (GSM409307) and H3K4me3 (GSM409308), MeRIP-seq (GSM1272365, GSM1272366, GSM1272367, GSM1272368). Additional materials, data, code, and associated protocols are available upon request.

## ACKNOWLEDGEMENTS

We thank Drs. Brigid Hogan (Duke) and Kenneth Poss (Duke) for feedback on previous versions of the manuscript, and Drs. Scott Soderling (Duke) and Christopher Nicchitta (Duke) for advising the BioID proteomics experiments. This work is supported by a new lab startup fund from Duke University Regeneration Next Initiative (current Duke Regeneration Center) (to Y.D.), Duke Whitehead Scholarship (to Y.D.), Glenn Foundation for Medical Research and AFAR Grants for Junior Faculty (to Y.D.), NIH 4D Nucleome Consortium U01HL156064 (to Y.D.), and NIH Genomic Innovator Awards R35HG011328 (to Y.D.). Y. Xiang is supported by a Duke Regeneration Center postdoctoral fellowship. Y. Xu is supported by Center for Advanced Genomic Technologies postdoctoral fellowship and the Duke Regeneration Center Fellowship to Accelerate Career Independence.

## AUTHOR CONTRIBUTIONS

T.S. and Y.D. conceived the idea of this study. T.S. performed all the experiments and carried out bioinformatics analysis with the help of Y.Xiang and Y.Xu. T.S. and E.S. performed LC-MS/MS proteomics analysis. T.S. and Y.D. wrote the paper.

## MATERIALS AND METHODS

### Cell culture

Human pluripotent stem cells are maintained in the chemically defined feeder-free, serum-free mTeSR plus medium (Stemcell Tech, 100-0276) according to manufacturer’s instructions and detached using 5mM EDTA/PBS or accutase. 10 μM Rho-associated protein kinase (ROCK) inhibitor Y-27632 was added into the culture medium if the cells were single cells. HEK293T cells were cultured in Dulbecco’s modified Eagle’s Medium (DMEM) supplemented with 10% FBS and 1% penicillin-streptomycin at 37□°C with 5% CO_2_.

### Embryoid body differentiation

To do spontaneous EB differentiation assay, 80% confluent H1 cells were dissociated using 496 5mM EDTA/PBS at room temperature for 10 min, and cells were aggregated in EB medium 497 (knockout DMEM/F12 supplemented with 20% knockout FBS, 1□× GlutaMax, 1□×_MEM 498 Non-Essential Amino Acids, 0.1 mM β-mercaptoethanol, and 1% penicillin-streptomycin) in 499 six-well ultra low attachment plates (Corning)^81^. Medium was changed every day.

### Neural cell differentiation

Neural cells were induced using STEMdiff™ SMADi Neural Induction Kit (Stemcell Technologies, 08581) following monolayer cell culture protocol. In brief, human stem cells were dissociated into single cells using accutase. Viable cells were counted using Trypan Blue and a hemocytometer. 5 x 10^5^ cells were transferred to a single well of 24 well plate in complete neural induction medium with 10 μM Y-27632. Medium was changed daily.

### Plasmid construction

All plasmids were constructed using restriction enzyme-based cloning, Gibson Assembly or Golden Gate Cloning. We made the dCas9-TurboID construct by replacing the KRBA domain on Lenti-dCas9-KRAB-blast (Addgene, 89567) with TurboID sequence (Addgene, 107169) using restriction enzyme-based cloning. We created the CARGO construct using Golden Gate Cloning. In brief, each single gRNA expression fragment was PCR amplified with BsmB1 cutting sites at both ends and ligated with each other and backbones using BsmB1 and T4 DNA ligase. We made dCasRX-ALKBH5 and dCasRX-dALKBH5 constructs using Gibson Assembly. dCasRX fragments (Addgen, 109050), ALKBH5 fragments (Addgene, 134783) or dALKBH5 fragments (Addgene, 134784) and backbones (Addgene, 89567) were PCR amplified, purified and assembled using Gibson Assembly Master Mix (NEB, E2611L). To make lenti-YTHDC2 construct, the cas9 sequence (Addgene, 52962) was replaced with full length YTHDC2 (Horizon, MHS6278-213245943) using Gibson Assembly. The point mutations were introduced by PCR amplification and validated by sanger sequencing.

### Transient transfection, lentivirus production and generation of stable cell lines

We performed transient transfection on human pluripotent stem cells using FuGENE® HD Transfection Reagent (Promega, E2311). The cells were splitted and treated with Y-27632 for one before transfection.

For lentivirus production, HEK293T cells were seeded one day before transfection. The next day, the media were changed with fresh medium and incubated for 2 hours. Plasmids and packaging plasmids psPAX2 and pMD.2G were co-transfected into cells using Lipofectamine™ 3000 Transfection Reagent (Invitrogen, L3000015). The medium was changed after 12 hour incubation. Viruses can be harvested at 36h, 60h and 84h post-transfection. Viruses were filtered through a 0.45 μm PES filter and concentrated using centrifugal filters (Genesee scientific, 84- μ 568).

To generate stable H1 cells, concentrated viruses were added into the medium with polybrene and Y-27632. The cells were spined at 1000g, 32°C for 90 min and then incubated in the incubator for another 2 hours before changing medium.

### Proximity labeling assay and Quantitative LC–MS/MS analysis

Stable cells expressing dCas9-TurboID and gRNAs were plated. When cells were 80% confluent the next day, biotin (Sigma, B4501) was added into the medium to the final concentration of 500 μM. To identify the optimal biotinylation time, the cells were incubated with biotin for 0, 1, 2, 3, 4 hours in the incubator^34^. The cells were washed with ice cold PBS and resuspended in 1 × L L Hypotonic Buffer (20 mM Tris-HCl, pH 7.4, 10 mM NaCl, 3 mM MgCl^2^) by pipetting up and down several times. After incubation on ice for 15 minutes, 25 μl 10% NP-40 were added and vortexed for 10 seconds. The lysates were spined for 15 minutes at 2000 g at 4°C. The supernatants were discarded. After washing with ice cold PBS twice, the pellets were resuspended in RIPA buffer (50 mM Tris-HCl, pH 8.0, 150 mM sodium chloride,1.0% Igepal CA-630, 0.5% sodium deoxycholate, and 0.1% sodium dodecyl sulfate) and sonicated to disrupt the nucleus. The nuclear lysates were by centrifugation at 12,000 g for 10 minutes at 4 °C to pellet the debris. The supernatants containing the biotinylated protein were transferred to a tube on ice for western blot or immunoprecipitation analysis. For western blot analysis, the same amount of proteins were loaded onto gradual NuPAGE Mini Protein Gels (Invitrogen, NP0323BOX). For immunoprecipitation analysis, 100 μl of streptavidin beads were washed twice with the RIPA buffer and added to samples. After incubation at 4°C overnight, the beads were washed twice with RIPA buffer and three times with IP wash buffer (50 mM Tris HCl pH 7.4, 150 mM NaCl, 0.1% SDS). The protein complexes were eluted by boiling in the 1 × L loading buffer at 95 °C for 10 min. The eluted proteins were loaded onto SDS-PAGE gel to analyze biotinylation efficiency.

For quantitative LC–MS/MS analysis, four dishes of hESCs were used for each replicate. Cells were treated with 500 μM biotin in the incubator. Cells were washed twice with ice cold PBS and transferred to 15 ml tubes. Cell pellets were resuspended in the IP lysis buffer (50 mM Tris HCl pH 7.4, 250 mM NaCl, 0.1 % SDS, 0.25 mM DTT) by pipetting up and down. The insoluble fractions were discarded after centrifugation. 200 μl magnetic streptavidin beads (Invitrogen, 65002) were washed three times in the IP buffer. The supernatants were incubated with washed beads at 4 °C overnight under gentle rotation. Beads were collected on the magnetic stand and washed five times stringently with the SDS buffer (50 mM Tris HCl pH 7.4, 150 mM NaCl, 0.1% SDS). After the final wash, biotinylated protein complexes were eluted with the 100 μl of elution buffer (2% SDS, 25 mM Tris pH 7.4, 5 mM DTT, and 5 mM free biotin) at 95°C for 10 min. The samples were loaded onto SDS-PAGE gel to run western blot and check IP efficiency vs negative control. The left samples in the elution buffer were delivered to the Duke proteomics facility for quantitative LC–MS/MS analysis.

### Generation of knockout cell lines

Guide RNAs(sgRNAs) targeting YTHDC2 were selected from Human CRISPR Knockout Pooled Library (GeCKO v2)^82^. YTHDC2 targeting gRNAs were cloned into lentiCRISPRv2 vectors and lentiviruses were packaged. hESCs were infected with viruses and selected with puromycin in six well plates. 1000 cells were transferred into 10 cm dishes. Single colonies were manually picked and replated individually into 24-well plates. Knockout efficiencies were detected by western blot. The genomic DNA of knockout cells were extracted and sequenced to validate genotypes.

### m^6^A editing by dCasRx-ALKBH5

To remove specific m^6^A sites on RNA, cells were co-transfected with dCasR-ALKBH5 and gRNAs targeting m^6^A peak region. dCasRx-dALKBH5 and scramble gRNA served as negative control. Cells were selected with puromycin or blastomycin. Samples were collected 5-7 days after transfection. Total RNA was extracted and used for real time quantitative PCR and m^6^A RIP-qPCR assays to detect RNA change and m^6^A level. The m^6^A peak negative region of edited RNA served as internal control for m^6^A level normalization

### Co-immunoprecipitation and western blot

Co-IP assay was performed as described previously^83^. We used four 15-cm dishes of hESCs to detect the interaction between YTHDC2 and TET1. hESCs were washed twice with PBS. Cells were crosslinked in 200μg/ml DSP/DMSO/PBS solution at room temperature for 20 min, then washed twice with PBS. Next, crosslink was stopped in 0.125mM glycine/PBS solution for 5 min. Cells were washed twice and collected in ice-cold PBS. Cell lysates were prepared in the Co-IP lysis buffer (50mM Tris-HCl pH7.5, 120mM NaCl, 0.5% NP-40, 0.1% SDS, 1 × EDTA L free protease inhibitor cocktail). The soluble fractions were collected after centrifugation at 12000 g for 15 min). The supernatants were transferred to new tubes and incubated with 5 μg anti-lgG (Cell Signaling Technology, 3900) or anti-YTHDC2 antibody (Abcam, ab220160) overnight on a rotator. Next day, 200 μl washed magnetic beads (Thermo Fisher Scientific, 11203D) were added into each sample. After 3 hours of incubation, beads were collected and washed 5 times with wash buffer (2mM Tris-HCl pH 8.0, 100mM NaCl, 1mM EDTA and 0.5% SDS loading buffer and analyzed by western blot.

### RNA extraction and Real-time Quantitative Analysis

We performed RNA extraction using Direct-zol RNA MiniPrep Kit (Zymo Research, 11-331) according to the manufacturer’s instructions. All RNA were treated with DNase during extraction to remove genomic DNA. RNA was reverse transcribed using SMARTScribe Reverse Transcriptase kit (TaKaRa, 639538). cDNA was diluted according to the RNA amount. Real-time Quantitative Analysis (RT-qPCR) was performed using iTaq Universal SYBR Green Supermix (Bio-Rad, 1725122). The results were computed using the 2^−ΔΔCT^method.

### RNA immunoprecipitation

RNA immunoprecipitation (RIP) was performed as previously reported^84^. For each replicate, two dishes of hESCs were washed with ice-cold PBS and transferred to a 15 ml tube. Cells were lysed with 10 ml lysis buffer (50 mM Tris-HCl pH 7.4, 150 mM NaCl, 1 mM DTT, 0.5% Igepal CA-630, supplemented with 40 U/ml of RNase inhibitor (Invitrogen, 10777019) and 1 × EDTA free protease inhibitor). After incubation at 4 °C for 30 min on a rotator, the lysates were spined at 12000 g for 15 min. The supernatants were transferred to a new 15 ml tube. 100 μ1 of supernatant were saved as input. The remaining lysates were incubated with 5 μg antibody (Abcam, ab220160) overnight on a rotator. Next day, 200 μ1 washed magnetic beads (Thermo Fisher Scientific, 11203D) were added into each sample. After 4 hours of incubation, beads were collected and washed 5 times with RIP lysis buffer at 4°C. The RNA complexes were eluted with 500 μl TRIzol reagent (Ambion, 15596108). The RNA was isolated using Direct-zol RNA MiniPrep Kit (Zymo Research, 11-331) and then subjected to RNA sequencing analysis.

### m^6^A-RIP

m^6^A-RIP was performed as previously reported^68, 85^. Total RNA was prepared using Direct-zol RNA MiniPrep Kit (Zymo Research, 11-331). DNA was digested during RNA preparation. For g of total RNA were fragmented into ∼200-nt-long fragments in 1 × RNA fragmentation buffer (10 mM Tris-HCl, 10 mM ZnCl^2^ in nuclease-free H2O) in a preheated thermal cycler for 5 minutes at 95°C. RNA were purified using RNA Clean & Concentrator-5 Kit (Zymo Research, R1014). Purified RNA were diluted with 1 × IP buffer (150 mM NaCl, 10 mM Tris-HCl pH 7.5, 0.1% Igepal CA-630 in nuclease-free H^2^O supplemented with RNase inhibitor). 15 μl anti-m^6^A antibodies (Cell Signaling Technology, D9D9W) were added and incubated at 4 °C overnight. 100 μl of protein-A magnetic beads (Thermo Fisher Scientific, 10002D) and 100 μl of protein-G m^6^A-RIP was performed as previously reported^68, 85^. Total RNA was prepared using Direct-zol RNA MiniPrep Kit (Zymo Research, 11-331). DNA was digested during RNA preparation. For each replicate, 10 μg of total RNA were fragmented into ∼200-nt-long fragments in 1□× RNA fragmentation buffer (10 mM Tris-HCl, 10 mM ZnCl^2^ in nuclease-free H2O) in a preheated thermal cycler for 5 minutes at 95°C. RNA were purified using RNA Clean & Concentrator-5 Kit (Zymo Research, R1014). Purified RNA were diluted with 1□× IP buffer (150 mM NaCl, 10 mM Tris-HCl pH 7.5, 0.1% Igepal CA-630 in nuclease-free H^2^O supplemented with RNase inhibitor). 15 μl anti-m^6^A antibodies (Cell Signaling Technology, D9D9W) were added and incubated at 4 °C overnight. 100 μl of protein-A magnetic beads (Thermo Fisher Scientific, 10002D) and 100 μl of protein-G magnetic beads (Thermo Fisher Scientific, 10004D) were washed three times with 1□× IP buffer and added into samples for another 4 hour incubation. The RNA-beads complexes were collected and washed 5 times using 1 mL of 1□×_IP buffer. The RNA complexes were eluted with 500 μl TRIzol reagent (Ambion, 15596108). The RNA was isolated using Direct-zol RNA MiniPrep Kit (Zymo Research, 11-331). Reverse transcription and real-time quantitative PCR analysis were used to measure m^6^A level.

### METTL3 inhibition and RNA stability assay

For METTL3 inhibition, STM2457 (MCE, HY-134836) were 646 added into the medium at final concentration of 5 μM for 1, 2, 3 days^64^. Total RNA was extracted for analysis of the m^6^A level of HERV-H. The cells were also used for HEVRH stability analysis. For RNA stability assay, cells were plated 16-24 hours before treatment. 80% confluent hESCs were treated with 5 μg ml^-1^ actinomycin D (Sigma, A1410) for 0, 1h, 3h, 7h to inhibit transcription. Total RNA was extracted and used for RT-qPCR analysis.

### Nascent RNA synthesis measured by RT-qPCR

We performed nascent RNA capture using Click-iT Nascent RNA Capture Kit (Thermo Fisher Scientific, C10365) according to the manufacturer’s instructions. Cells were plated on matrigel coated six well plates. When cells were 80% confluency, 5-Ethynyl Uridine (EU) was added to 0.5 mM for 60 min and 10 min in the incubator. Total RNA was extracted using Direct-zol RNA MiniPrep Kit. EU labeled RNA was biotinylated and purified using streptavidin beads. The enriched RNA was reverse transcribed and used for RT-qPCR analysis.

### RNA sequencing and data analysis

Libraries for RNA sequencing were generated using Smart-seq2^86^. Total RNA was extracted using Direct-zol RNA MiniPrep Kit following the manufacturer’s instructions. RNA concentrations were measured on the Qubit. 10 ng of total RNA was used for library construction. RNA was reverse transcribed. cDNA was used to generate double stranded DNA (dsDNA). dsDNA were purified using magnetic beads. 30 ng of purified dsDNA was tagmented at 55 °C for 15 min and purified using DNA Clean & Concentrator-5 kit (Zymo Research, D4004) The tagmented DNA was PCR amplified using Nextera primers. The libraries were purified using magnetic beads and sequenced using the MGISEQ-2000 in pair-end mode with 100 base pair (bp) per read.

For RNA-seq data analysis, raw reads were trimmed using Fastp (0.20.0)^87^ and aligned to the hg38 genome using HISAT2 (2.1.0)^88^ with default settings. Reads on each gene were counted using featureCounts (2.0.1)^89^. Differentially expressed genes were determined using DESeq2 (1.36.0)^90^. Gene ontology analysis was performed using the DAVID webtool^91, 92^. Overlapped genes were performed using the Calculate and draw custom Venn diagrams webtool (http://bioinformatics.psb.ugent.be/webtools/Venn/).

### ChIP-seq and data analysis

H3K9me3 and H3K27ac ChIP assays were performed as reported previously^34^. Cells were crosslinked with the crosslinking buffer (10mM NaCl, 0.1 mM EDTA, 0.05 mM EGTA, 5 mM Hepes pH8.0, 1% formaldehyde) at room temperature for 10 min. The crosslinking reaction was stopped by adding 2.5M glycine to a final concentration of 0.125M and incubation at room temperature for another 5 min. Cells were washed with ice cold PBS twice and transferred to 15 mL tubes in PBS. After centrifugation at 500g for 5 min, the PBS was discarded. Cell pellets were resuspended in 50 μl lysis buffer (1% SDS, 50 mM Tris-HCl, pH 8.0, 20 mM EDTA supplemented with 10x protease inhibitor) and incubated for10 min on ice. The cell lysates were diluted to 500 μl with the cold TE buffer. The lysates were sonicated using Covaris M220. The sonicated lysates were centrifuged at 12000 g for 15 min at 4 °C. The supernatants were transferred into new tubes and DNA concentrations were measured on a NanoDrop. For each g of chromatin was used. To pre-clear the chromatin solutions, 20 μl 1gG Dynabeads (Life Technologies, 11204D) were washed and added. The chromatin beads solutions were rotated gently in the cold room for 2 hours. After the incubation, place tubes on a magnetic stand to collect beads. 100 μl supernatants were transferred into new tubes. 100 μl Binding Buffer (1% Triton-X, 01% Sodium Deoxycholate supplemented with protease inhibitor in TE buffer) was added to the 100 μl chromatin. The mixture was incubated in the cold room overnight on a rotator. Next day, 20 μl lgG pre-washed Dynabeads were added into each reaction and incubated for another 2 hours. After incubation, the beads were washed five times with wash buffer (50 mM Hepes, pH 8.0, 1% NP-40, 0.70% Sodium Deoxycholate, 0.5 M LiCl, 1 mM EDTA supplemented with protease inhibitor) and twice with cold 1 × TE. After the final wash, 150 μl ChIP elution buffer (10 mM, Tris-HCl pH 8.0, 1 mM EDTA, 1% SDS) were added into tubes. The beads mixture was transferred to new tubes and incubated at 65°C for 20 minutes at 1300 rpm on a Thermomixer. The supernatants were transferred into tubes and incubated at × TE and 8 μl of 10 mg/mL RNase A μl of 20 mg/mL Proteinase K was was added to each sample and incubated at 37°C for 1 hour. 8 μl of 20mg/mL Proteinase K was added and incubated at 55°C for 1 hour. After that, the DNA was purified and measured on a NanoDrop. To generate ChIP libraries, DNA ends were repaired. ployA tails were added to the ends and then adaptors were ligated at room temperature. Last, the DNA was amplified with TruSeq primers and purified with magnetic beads. Libraries were sequenced on the MGISEQ-2000 platform in pair-end mode with 100 base pair (bp) per read.

For H3K9me3 and H3K27ac ChIP-seq analyses, raw reads were trimmed using Fastp (0.20.0)^87^ and aligned to the hg38 genome using Bowtie2 (2.4.4)^93^ with the parameter ‘bowtie2 -X 2000 -x -1 -2 --fr --rg-id --rg DS: --rg PL: --rg SM: -p 24’. For TET1 and YTHDC2 ChIP-seq analysis, reads were trimmed using Trim Galore (0.6.4) (https://www.bioinformatics.babraham.ac.uk/projects/trim_galore/) and aligned to the hg38 genome using BWA (0.7.17)^94^. Mapped reads were filtered by MAPQ > 10 using SAMtools (1.12)^95^. PCR duplicates were found using Sambamba (0.8.0)^96^ with the options ‘sambamba markdup -t 10 --sort-buffer-size=6000 --overflow-list-size 600000’. Peak calling was performed using MACS2 (2.2.7.1)^97^ with the following options ‘macs2 callpeak -t -c -f BAM -g 2945849067 --qvalue 0.001 --keep-dup all --outdir --name --nomodel --mfold 0 50’. HOMER (4.10)^98^ annotatePeaks.pl was used to annotate peaks and gene ontology analysis. Genome coverage bigWig files for visualization were generated through bamCoverage with the options ‘bamCoverage -b -o --binSize 25 -p 16 --normalizeUsing RPKM’ using deeptools (3.5.0)^99^. Heatmaps and profiles plots were generated using deeptools computerMatrix and plotProfile. The overlapping peaks were calculated using bedtools intersect or HOMER mergePeaks. The Jaccard statistic ratio of the intersection of two sets of peaks was calculated using bedtools (2.30.0)^100^.

### CUT&RUN and data analysis

We performed CUT&RUN using CUTANA™ ChIC/CUT&RUN Kit (EpiCypher, 14-1048) according to manufacturer’s instructions. For each replicate, 11 μl ConA Beads were activated in the Bead Activation Buffer and kept on ice until cells were ready. For BRD4 CUT&RUN assay, 200 000 cells were used and slightly crosslinked in 0.1% formaldehyde for 2 min at room temperature. Cells were washed twice using PBS and transferred to tubes. Cells were washed twice in 100 μl Wash Buffer . After final wash, cells were resuspended in 100 μl Wash Buffer. 10 μl pre-washed ConA Beads were added to cells, mixed by pipetting up and down 5-10 times gently and transferred to PCR tubes. After incubation for 10 min at room temperature, cells were resuspended in the 50 μl ice cold Antibody Buffer. 1 μl antibodies were added and incubated in the cold room overnight on a rotator. Then beads were washed twice in 200 μl cold Cell Permeabilization Buffer and completely resuspended in 50 μl cold Cell Permeabilization Buffer. 2.5 μl pAG-MNase was added and incubated on the rotator for 10 min at room temperature. Then beads were washed twice in 200 μl cold Cell Permeabilization Buffer and completely resuspended in 50 μl cold Cell Permeabilization Buffer. 1 μl 100 mM Calcium Chloride was added to each tube and incubated for two hours in the cold room. At the end of incubation, 33 μl Stop Buffer Master Mix was added to each reaction and incubated for 30 min at 37 °C to release CUT&RUN fragments from the insoluble nuclear chromatin. DNA containing supernatants were transferred to new 1.5 mL tubes and DNA was purified and used for library preparation. Libraries were sequenced on the MGISEQ-2000 platform in pair-end mode with 100 bp per read.

For CUT&RUN data analysis, raw reads were trimmed using Fastp (0.20.0)^87^ and aligned to the hg38 genome using Bowtie2 (2.4.4)^93^. Mapped reads were filtered by MAPQ > 10 using SAMtools (1.12)^95^. PCR duplicates were found using Sambamba (0.8.0)^96^. Peak calling was performed using MACS2 (2.2.7.1)^97^.

### MeDIP sequencing and data analysis

Genomic DNA from YTHDC2 WT or YTHDC2 knockout cells were diluted in TE buffer (10 mM Tris-HCl pH8.0, 1 mM EDTA) and sonicated using Covaris. The sonicated DNA was purified with DNA Clean & Concentrator-5 kit (Zymo Research, D4004). 10 μg purified DNA was repaired, added A bases to the 3′ end and ligated with PCR adapters. The DNA was purified using the kit again. 1 μg DNA in 450 μl TE buffer was denatured by boiling at 95°C for 10 min, followed by snap chilling on ice. 50 μl 10x IP buffer (1.4 M NaCl, 100 mM Na-phosphate, pH 7.0, 0.5% Triton X-100) was added to the DNA solution and mixed by pipetting up and down 5-10 times. 1 μl anti-5hmC or anti-5mC antibodies were added and incubated i n the cold room overnight. 100 μl magnetic beads were washed in the 1□×_IP buffer and added. After incubation at 4°C for another 4 hours, the beads were collected and washed five times in the 1□×_IP buffer. DNA was eluted from beads by boiling at 65°C for 15 min in the elution buffer (10 mM Tris pH 7.6, 1 mM EDTA, 1% SDS). The DNA was purified and PCR amplified. Libraries were size selected and sequenced on the MGISEQ-2000 platform in pair-end mode with 100 bp per read.

For MeDIP-seq data analysis, raw reads were trimmed using Fastp (0.20.0)^87^ and aligned to the hg38 genome using Bowtie2 (2.4.4)^93^. Mapped reads were filtered by MAPQ > 30 using SAMtools (1.12)^95^. PCR duplicates were found using Sambamba (0.8.0)^96^. Peak calling was performed using MACS2 (2.2.7.1)^97^.

### ATAC sequencing and data analysis

For each replicate, 50000 cells were lysed in NPB buffer (PBS containing 5% BSA, 1mM DTT, 0.2% IGEPAL, 1□×_Roche Complete Protease Inhibitor) and incubated on ice for 15 min to isolate nuclei. After centrifugation, the nuclei were resuspended in 100 μl 1□× TB buffer (33mM Tris-AC pH 7.8, 66mM K-AC, 10mM Mg-AC, 16% DMF). 2 μl homemade Tn5 transposase were added to each reaction. Nuclei were then incubated 37 °C for 1 hour on a thermomixer at 1000 rpm. After incubation, genomic DNA was purified using DNA Clean & Concentrator-5 kit and dissolved in 15 μl DNase free H^2^O. The DNA was PCR amplified using Nextera primers. Libraries were size selected and sequenced on the MGISEQ-2000 platform in pair-end mode with 100 bp per read.

For ATAC-seq data analysis, raw reads were trimmed using Fastp (0.20.0)^87^ and aligned to the hg38 genome using Bowtie2 (2.4.4)^93^. Mapped reads were filtered by MAPQ > 30 using SAMtools (1.12)^95^. PCR duplicates were found using Sambamba (0.8.0)^96^. Peak calling was performed using MACS2 (2.2.7.1)^97^.

## EXTENDED DATA FIGURES AND LEGENDS

### Figure legends for the Extended Data Figure. 1-7

**Extended Data Fig. 1.**
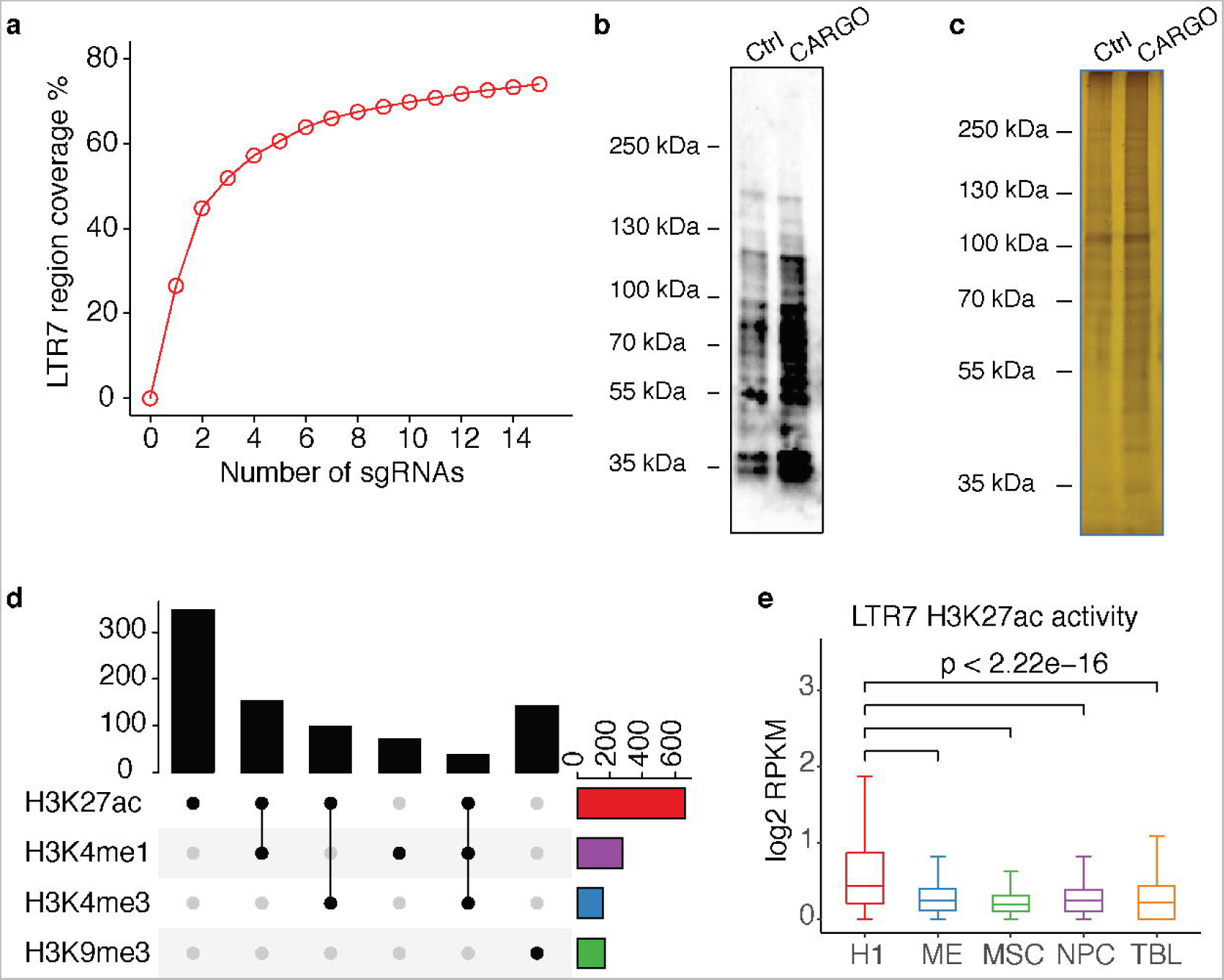
CARGO targets LTR7 sequences that are associated with active chromatin marks in human embryonic stem cells. **a,** A total of 15 sgRNAs are expressed from the CARGO construct. The sgRNAs target the conserved sequences of LTR7 loci. The predicted coverage ratio of LTR7 loci is calculated based on the sequence similarity of sgRNA and LTR7 sequences. **b, c,** The purified biotinylated nuclear proteins used for proteomics analysis were analyzed by western blot (**b**) and silver staining (**c**). **d,** Classification of LTR7 loci based on their histone modification signature in H1 hESCs. H1 hESC ChIP-seq datasets are downloaded from ENCODE data portal. **e,** Histone H3K27ac ChIP-seq signals of LTR7 loci in H1 hESCs and H1-derived differentiated lineages, including mesendoderm (ME), mesenchymal stem cell (MSC), neural progenitor cells (NPC), and trophoblast-like cells (TBL). The ChIP-seq datasets are generated by Epigenome Roadmap projects. P values were calculated by Wilcoxon signed-rank test.

**Extended Data Fig. 2.**
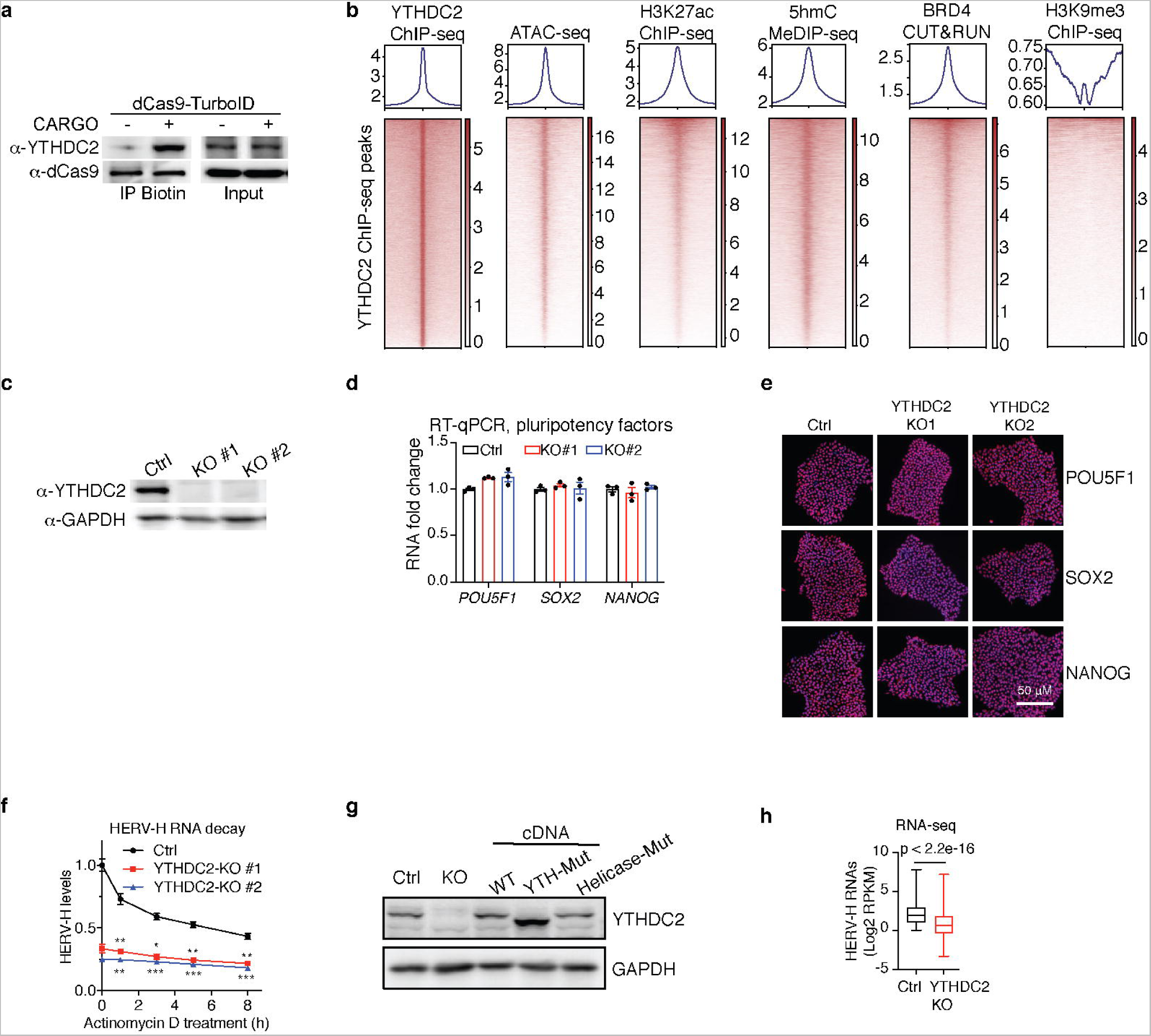
YTHDC2 occupies active chromatin loci and is dispensable for pluripotency maintenance of hESCs. **a,** The nuclear proteins prepared in the same way for CARGO-BioID proteome analysis were immunoprecipitated by Streptavidin beads followed by western blot analysis, to validate the spatial proximity of YTHDC2 to LTR7 genomic loci. **b,** The indicated ChIP-seq, ATAC-seq, MeDIP-seq and CUT&RUN data are presented as heatmaps centered on YTHDC2 peaks. Each row of the heatmap represents a YTHDC2 peak. **c,** Detection of YTHDC2 knockout efficiency in the two hESC clones using western blot. **d,** RT-qPCR detection of pluripotency markers expression in wild-type (WT) control versus YTHDC2 KO hESCs. **e,** Immunofluorescence staining showing pluripotency markers in WT and two independent YTHDC2 KO clones. **f,** RT-qPCR detection of HERV-H transcripts decay rate in WT and YTHDC2 KO cells treated with Actinomycin D, which block new RNA synthesis. **g,** Western blot detection of YTHDC2 knockout efficiency and YTHDC2 overexpression level in hESCs. WT indicates WT full-length YTHDC2 cDNA. YTH-Mut indicates YTHDC2 cDNA with YTH domain deletion. Helicase-Mut indicates YTHDC2 cDNA with helicase domain mutation. **h,** Boxplot showing expression level of HERV-H in WT and YTHDC2 KO cells detected by RNA-seq. Data represent mean ± SEM from three independent experiments. * p < 0.05, ** < p < 0.01, *** p < 0.001. P values were calculated by two-tailed Student’s test (**f**) and Wilcoxon signed-rank test (**h**).

**Extended Data Fig. 3.**
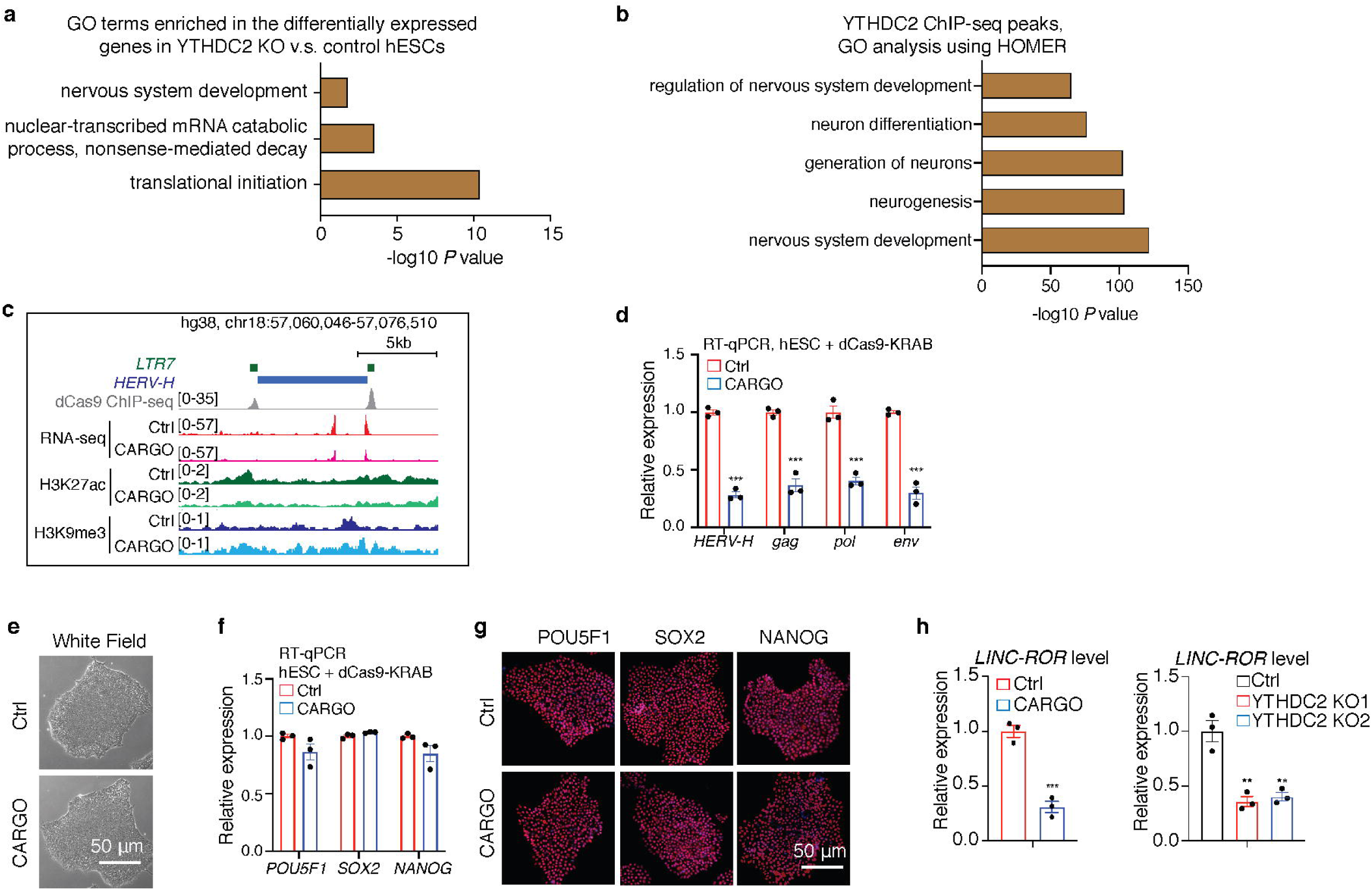
The YTHDC2/LTR7 axis is dispensable for pluripotency maintenance of hESCs in the self-renewal medium. **a,** GO analysis for differentially expressed genes in WT and YTHDC2 KO cells. RNA translation and decay terms were enriched, which is consistent with its m^6^A reader function. Enriched nervous system development term suggests its potential function on cell fate. **b,** GO analysis for YTHDC2 ChIP-seq peaks showing enriched neural development associated terms. **c,** UCSC genome browser snapshot showing dCas9 binding, RNAseq, H3K27ac and H3K9me3 ChIP-seq signal on LTR7/HERV-H in Ctrl and CARGO expressing hESCs. **d,** RT-qPCR showing markedly decrease of HERV-H transcripts and the HERV-H viral genes expression in Ctrl and LTR7-silenced hESCs. **e,** White field images showing morphologies of Ctrl and CARGO expressing H1 hESCs. **f,** RT-qPCR detection of indicated pluripotency markers in Ctrl versus CARGO expressing H1 hESCs. **g,** Immunofluorescence staining of indicated pluripotency markers in Ctrl versus CARGO expressing H1 hESCs. **h,** RT-qPCR detection of LINC-ROR in Ctrl and CARGO expressing H1 hESCs (left) and in YTHDC2 Ctrl and YTHDC2 KO H1 hESCs (right). Data represent mean ± SEM from three independent experiments. *p < 0.05, **p < 0.01, ***p < 0.001. P values were calculated by two-tailed Student’s test.

**Extended Data Fig. 4.**
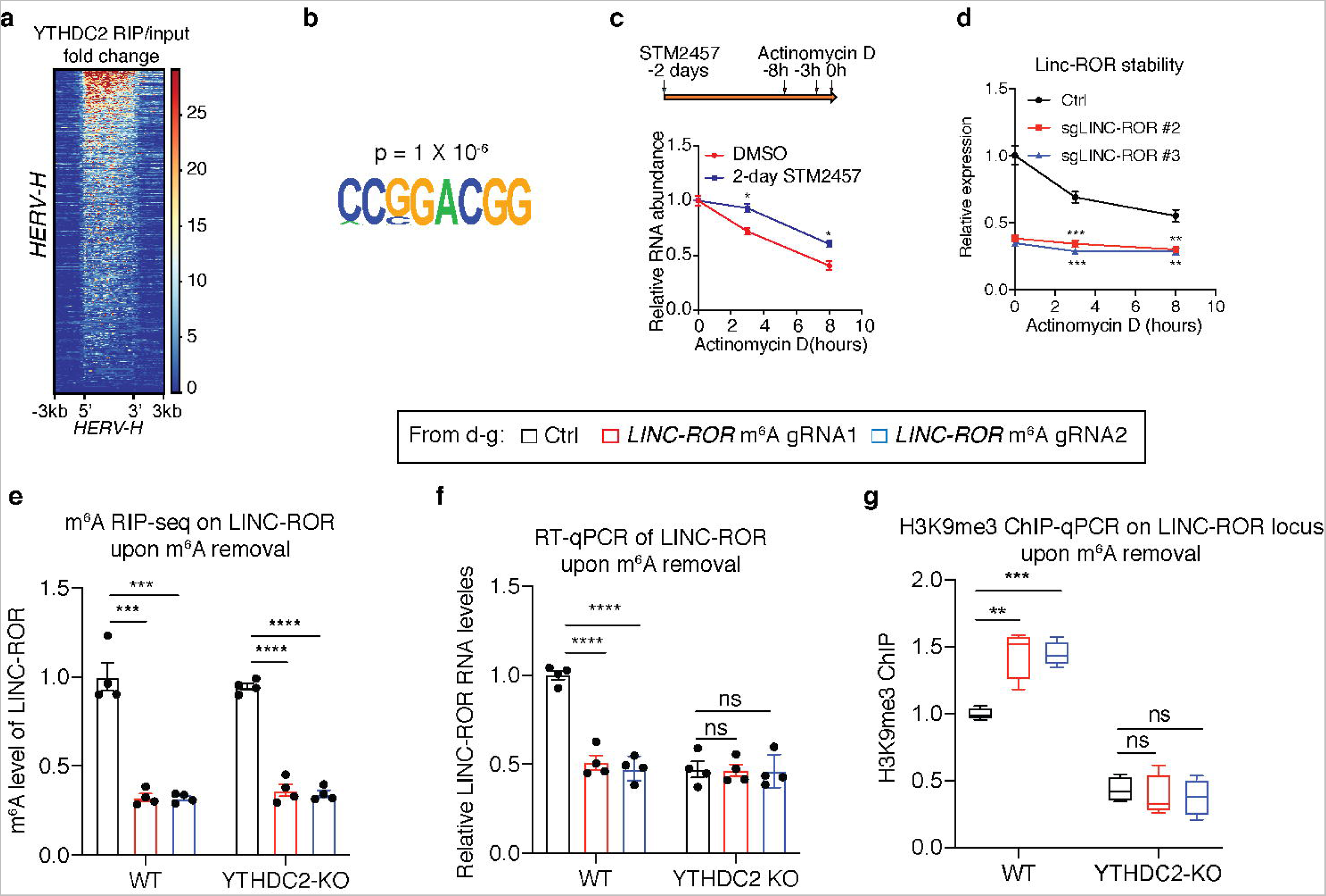
m^6^A-modified HERV-H RNAs recruit YTHDC2 to LTR7/HERV-H genomic loci in a m^6^A-depenedent manner. **a,** Heatmap showing YTHDC2 RIP-seq signal (normalized against IgG control) along HERV-H transcripts. **b,** Motif analysis for YTHDC2 RIP-seq peaks showing enriched GGAC motif. **c,** After STM2457 treatment, the new RNA synthesis is blocked by Actinomycin D, followed by RT-qPCR quantification of HERV-H decay rate in DMSO and STM2457-treated hESCs. **d,** Upon removal of LINC-ROR m^6^A, the new RNA synthesis is blocked by Actinomycin D. RT-qPCR quantification was conducted in a time course manner to determine the decay rate of LINC-ROR in hESCs. **e-g,** MeRIP-qPCR analysis for m^6^A level (**e**), RT-qPCR analysis of HERV-H RNA expression (**f**) and ChIP-qPCR detection of H3K9me3 signal on nearby LTR7 (**g**) in WT and YTHDC2 KO cells transduced with indicated gRNAs. Data represent mean ± SEM from three independent experiments. ns, not significant; *p < 0.05, **p < 0.01, ***p < 0.001. P values were calculated by two-tailed Student’s test.

**Extended Data Fig. 5.**
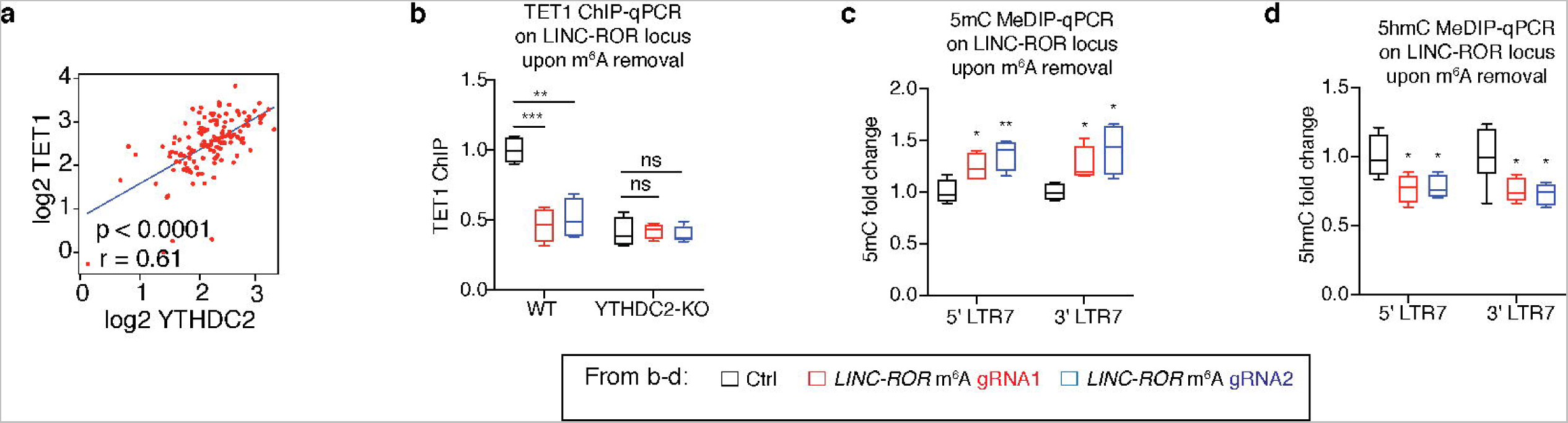
YTHDC2 recruits TET1 to demethylate LTR7/HERV-H DNA. **a,** Scatter plot showing correlation between YTHDC2 ChIP-seq signal and TET1 ChIP-seq signal on their co-occupied LTR7 loci in hESCs. **b,** ChIP-qPCR analysis of TET1 signal on LINC-ROR nearby LTR7 in WT and YTHDC2 KO cells transduced with indicated gRNAs. **c, d,** MeDIP-qPCR detection of 5mC (**c**) and 5hmC level (**d**) in H1 WT cells transduced with indicated gRNAs. Data represent mean ± SEM from three independent experiments. ns, not significant; *p < 0.05, **p < 0.01, ***p < 0.001. P values were calculated by two-tailed Student’s test.

**Extended Data Fig. 6.**
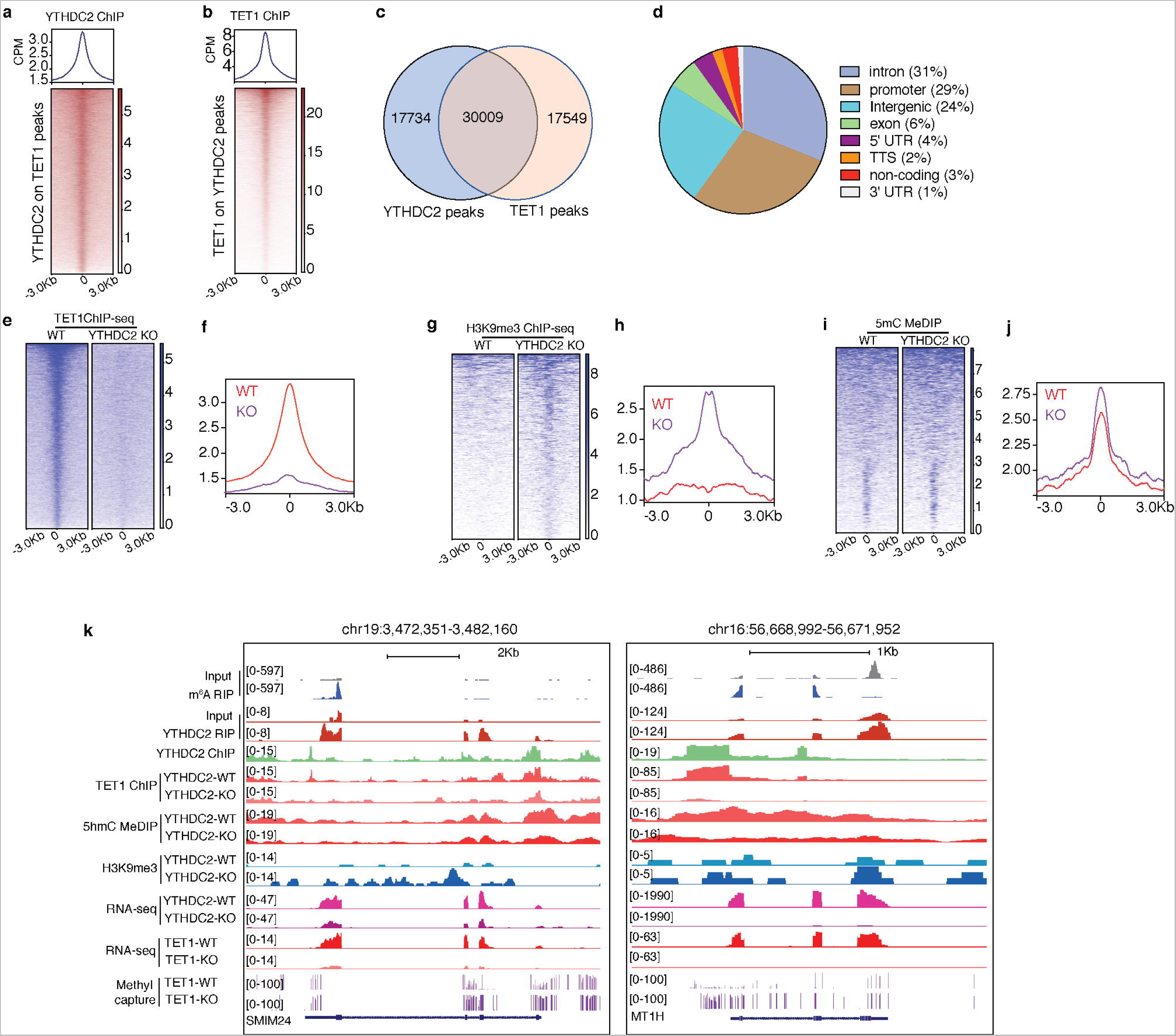
Crosstalk between YTHDC2 and TET1 across the genome. **a,** Heatmap (bottom) and aggregate signal plots (top) showing the endogenous YTHDC2 ChIP-seq signal centered on the peaks that are bound by TET1. **b,** Heatmap (bottom) and aggregate signal plot (top) showing TET1 ChIP-seq signal centered on the peaks that are bound by YTHDC2. **c,** Venn diagrams showing the overlap of YTHDC2 peaks and TET1 peaks. **d,** YTHDC2 and TET1 peaks categorized by chromatin feature. **e-j,** Heatmaps (**e**, **g** and **i**) and aggregate signal plots (**f**, **h** and **j**) showing TET1 ChIP-seq signal, H3K9me3 ChIP-seq signal and 5mC MeDIP-seq signal on YTHDC2 and TET1 co-bound peaks. **k,** Two representative UCSC genome browser snapshots illustrating features of YTHDC2 and TET1 co-bound genes and their transcripts.

**Extended Data Fig. 7.**
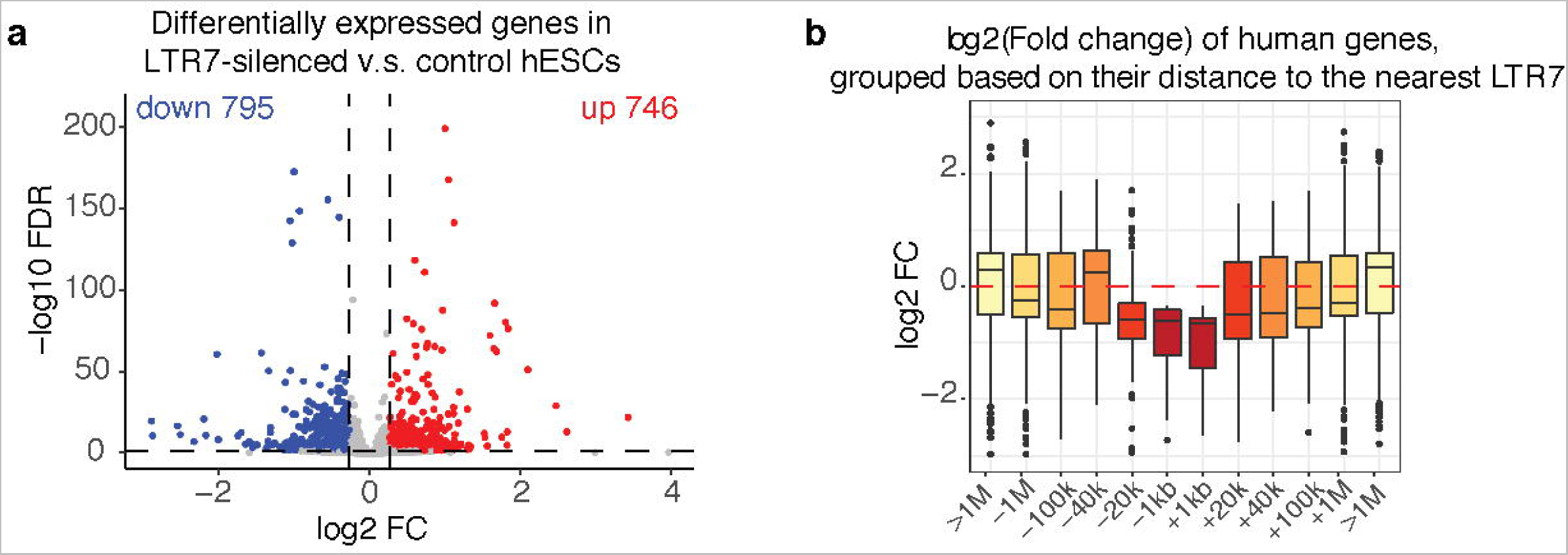
LTR7 sequences function as distal enhancers of human genes in hESCs. **a,** Volcano plot showing differentially expressed genes in Ctrl and CARGO expressing hESCs (DEseq2, fold change > 1.5, *P* < 0.05). There are 795 down-regulated (blue) and 746 upregulated genes in the LTR7-silenced hESCs. **b,** Box plot showing gene expression changes grouped by their distance to their nearest LTR7 in Ctrl and CARGO expressing hESCs. Gene expression changes are correlated with their distance to LTR7 upon LTR7 inhibition. LTR7 nearby genes show downregulation. LTR7 distal genes show upregulation.

## SUPPLEMENTARY TABLES

**Table 1. Oligo sequences used in this study.**

**Table 2. Results of CARGO-BioId proteomics analysis on LTR7s.**

**Table 3. Ingenuity Pathways Analysis analysis for proteins on LTR7s.**

**Table 4. Gene Ontology analysis for proteome.**

**Table 5. RNA-seq analysis of Ctrl versus LTR7 silenced H1 hESC cells.**

**Table 6. RNA-seq analysis of Ctrl versus YTHDC2-KO H1 hESC cells.**

**Table 7. RNA-seq analysis of wild-type H1 hESCs versus H1 hESC-derived neuronal cells.**

**Table 8. List of antibodies used in this study.**

## Notes

### Competing Interest Statement

The authors have declared no competing interest.

## REFERENCES

1. Bourque, G. et al. Ten things you should know about transposable elements. Genome Biol. 19, 199 (2018).

2. Chuong, E. B., Elde, N. C. & Feschotte, C. Regulatory activities of transposable elements: from conflicts to benefits. Nat. Rev. Genet. 18, 71–86 (2017).

3. Senft, A. D. & Macfarlan, T. S. Transposable elements shape the evolution of mammalian development. Nat. Rev. Genet. (2021) doi:10.1038/s41576-021-00385-1.

4. Fueyo, R., Judd, J., Feschotte, C. & Wysocka, J. Roles of transposable elements in the regulation of mammalian transcription. Nat. Rev. Mol. Cell Biol. 23, 481–497 (2022).

5. Deniz, Ö., Frost, J. M. & Branco, M. R. Regulation of transposable elements by DNA modifications. Nat. Rev. Genet. 20, 417–431 (2019).

6. Padeken, J., Methot, S. P. & Gasser, S. M. Establishment of H3K9-methylated heterochromatin and its functions in tissue differentiation and maintenance. Nat. Rev. Mol. Cell Biol. (2022) doi:10.1038/s41580-022-00483-w.

7. Geis, F. K. & Goff, S. P. Silencing and Transcriptional Regulation of Endogenous Retroviruses: An Overview. Viruses 12, (2020).

8. Maksakova, I. A. et al. H3K9me3-binding proteins are dispensable for SETDB1/H3K9me3-dependent retroviral silencing. Epigenetics Chromatin 4, 12 (2011).

9. Tchasovnikarova, I. A. et al. GENE SILENCING. Epigenetic silencing by the HUSH complex mediates position-effect variegation in human cells. Science 348, 1481–1485 (2015).

10. Cosby, R. L., Chang, N.-C. & Feschotte, C. Host-transposon interactions: conflict, cooperation, and cooption. Genes Dev. 33, 1098–1116 (2019).

11. Hermant, C. & Torres-Padilla, M.-E. TFs for TEs: the transcription factor repertoire of mammalian transposable elements. Genes Dev. 35, 22–39 (2021).

12. Hermann, A., Goyal, R. & Jeltsch, A. The Dnmt1 DNA-(cytosine-C5)-methyltransferase methylates DNA processively with high preference for hemimethylated target sites. J. Biol. Chem. 279, 48350–48359 (2004).

13. Okano, M., Bell, D. W., Haber, D. A. & Li, E. DNA methyltransferases Dnmt3a and Dnmt3b are essential for de novo methylation and mammalian development. Cell 99, 247– 257 (1999).

14. Ito, S. et al. Role of Tet proteins in 5mC to 5hmC conversion, ES-cell self-renewal and inner cell mass specification. Nature 466, 1129–1133 (2010).

15. Tahiliani, M. et al. Conversion of 5-methylcytosine to 5-hydroxymethylcytosine in mammalian DNA by MLL partner TET1. Science 324, 930–935 (2009).

16. Liu, J. et al. N6-methyladenosine of chromosome-associated regulatory RNA regulates chromatin state and transcription. Science 367, 580–586 (2020).

17. Xu, W. et al. METTL3 regulates heterochromatin in mouse embryonic stem cells. Nature (2021) doi:10.1038/s41586-021-03210-1.

18. Liu, J. et al. The RNA m6A reader YTHDC1 silences retrotransposons and guards ES cell identity. Nature 591, 322–326 (2021).

19. Wei, J. et al. FTO mediates LINE1 m6A demethylation and chromatin regulation in mESCs and mouse development. Science eabe9582 (2022).

20. Li, Y. et al. N6-Methyladenosine co-transcriptionally directs the demethylation of histone H3K9me2. Nat. Genet. 52, 870–877 (2020).

21. Lee, J.-H. et al. Enhancer RNA m6A methylation facilitates transcriptional condensate formation and gene activation. Mol. Cell 81, 3368–3385.e9 (2021).

22. Shi, H., Wei, J. & He, C. Where, When, and How: Context-Dependent Functions of RNA Methylation Writers, Readers, and Erasers. Mol. Cell 74, 640–650 (2019).

23. Yang, Y., Hsu, P. J., Chen, Y.-S. & Yang, Y.-G. Dynamic transcriptomic m6A decoration: writers, erasers, readers and functions in RNA metabolism. Cell Res. 28, 616–624 (2018).

24. Michalak, E. M., Burr, M. L., Bannister, A. J. & Dawson, M. A. The roles of DNA, RNA and histone methylation in ageing and cancer. Nat. Rev. Mol. Cell Biol. 20, 573–589 (2019).

25. Zhao, S., Allis, C. D. & Wang, G. G. The language of chromatin modification in human cancers. Nat. Rev. Cancer 21, 413–430 (2021).

26. Jambhekar, A., Dhall, A. & Shi, Y. Roles and regulation of histone methylation in animal development. Nat. Rev. Mol. Cell Biol. 20, 625–641 (2019).

27. Parry, A., Rulands, S. & Reik, W. Active turnover of DNA methylation during cell fate decisions. Nat. Rev. Genet. 22, 59–66 (2021).

28. Greenberg, M. V. C. & Bourc’his, D. The diverse roles of DNA methylation in mammalian development and disease. Nat. Rev. Mol. Cell Biol. 20, 590–607 (2019).

29. Erdmann, R. M. & Picard, C. L. RNA-directed DNA Methylation. PLoS Genet. 16, e1009034 (2020).

30. Matzke, M. A. & Mosher, R. A. RNA-directed DNA methylation: an epigenetic pathway of increasing complexity. Nat. Rev. Genet. 15, 394–408 (2014).

31. Hsu, P. J. et al. Ythdc2 is an N6-methyladenosine binding protein that regulates mammalian spermatogenesis. Cell Res. 27, 1115–1127 (2017).

32. Fuentes, D. R., Swigut, T. & Wysocka, J. Systematic perturbation of retroviral LTRs reveals widespread long-range effects on human gene regulation. Elife 7, (2018).

33. Gu, B. et al. Transcription-coupled changes in nuclear mobility of mammalian cis-regulatory elements. Science 359, 1050–1055 (2018).

34. Branon, T. C. et al. Efficient proximity labeling in living cells and organisms with TurboID. Nat. Biotechnol. 36, 880–887 (2018).

35. Kim, D. I. et al. Probing nuclear pore complex architecture with proximity-dependent biotinylation. Proc. Natl. Acad. Sci. U. S. A. 111, E2453–61 (2014).

36. Schmidtmann, E., Anton, T., Rombaut, P., Herzog, F. & Leonhardt, H. Determination of local chromatin composition by CasID. Nucleus 7, 476–484 (2016).

37. Gao, X. D., Rodríguez, T. C. & Sontheimer, E. J. Adapting dCas9-APEX2 for subnuclear proteomic profiling. Methods Enzymol. 616, 365–383 (2019).

38. Liu, X. et al. Multiplexed capture of spatial configuration and temporal dynamics of locus-specific 3D chromatin by biotinylated dCas9. Genome Biol. 21, 59 (2020).

39. Ugur, E., Bartoschek, M. D. & Leonhardt, H. Locus-Specific Chromatin Proteome Revealed by Mass Spectrometry-Based CasID. Methods Mol. Biol. 2175, 109–121 (2020).

40. Xie, W. et al. Base-resolution analyses of sequence and parent-of-origin dependent DNA methylation in the mouse genome. Cell 148, 816–831 (2012).

41. Leung, D. et al. Integrative analysis of haplotype-resolved epigenomes across human tissues. Nature 518, 350–354 (2015).

42. Zhang, Y. et al. Transcriptionally active HERV-H retrotransposons demarcate topologically associating domains in human pluripotent stem cells. Nat. Genet. 51, 1380–1388 (2019).

43. Glinsky, G. V. Transposable Elements and DNA Methylation Create in Embryonic Stem Cells Human-Specific Regulatory Sequences Associated with Distal Enhancers and Noncoding RNAs. Genome Biol. Evol. 7, 1432–1454 (2015).

44. Gorkin, D. U., Leung, D. & Ren, B. The 3D genome in transcriptional regulation and pluripotency. Cell Stem Cell 14, 762–775 (2014).

45. Shi, J. & Vakoc, C. R. The mechanisms behind the therapeutic activity of BET bromodomain inhibition. Mol. Cell 54, 728–736 (2014).

46. Dhalluin, C. et al. Structure and ligand of a histone acetyltransferase bromodomain. Nature 399, 491–496 (1999).

47. Murakami, S. & Jaffrey, S. R. Hidden codes in mRNA: Control of gene expression by m6A. Mol. Cell 82, 2236–2251 (2022).

48. He, P. C. & He, C. m6 A RNA methylation: from mechanisms to therapeutic potential. EMBO J. 40, e105977 (2021).

49. Batista, P. J. et al. m6A RNA Modification Controls Cell Fate Transition in Mammalian Embryonic Stem Cells. Cell Stem Cell 15, 707–719 (2014).

50. Jao, C. Y. & Salic, A. Exploring RNA transcription and turnover *in vivo* by using click chemistry. Proc. Natl. Acad. Sci. U. S. A. 105, 15779–15784 (2008).

51. Saito, Y. et al. YTHDC2 control of gametogenesis requires helicase activity but not m6A binding. Genes Dev. 36, 180–194 (2022).

52. Li, L. et al. The XRN1-regulated RNA helicase activity of YTHDC2 ensures mouse fertility independently of m6A recognition. Mol. Cell 82, 1678–1690.e12 (2022).

53. Larivera, S. & Meister, G. Domain confusion 2: m6A-independent role of YTHDC2. Molecular cell vol. 82 1608–1609 (2022).

54. Lu, X. et al. The retrovirus HERVH is a long noncoding RNA required for human embryonic stem cell identity. Nat. Struct. Mol. Biol. 21, 423–425 (2014).

55. Wang, J. et al. Primate-specific endogenous retrovirus-driven transcription defines naive-like stem cells. Nature 516, 405–409 (2014).

56. Ohnuki, M. et al. Dynamic regulation of human endogenous retroviruses mediates factor-induced reprogramming and differentiation potential. Proc. Natl. Acad. Sci. U. S. A. 111, 12426–12431 (2014).

57. Koyanagi-Aoi, M. et al. Differentiation-defective phenotypes revealed by large-scale analyses of human pluripotent stem cells. Proceedings of the National Academy of Sciences vol. 110 20569–20574 (2013).

58. Chambers, S. M. et al. Highly efficient neural conversion of human ES and iPS cells by dual inhibition of SMAD signaling. Nat. Biotechnol. 27, 275–280 (2009).

59. Thakore, P. I. et al. Highly specific epigenome editing by CRISPR-Cas9 repressors for silencing of distal regulatory elements. Nat. Methods 12, 1143–1149 (2015).

60. Loewer, S. et al. Large intergenic non-coding RNA-RoR modulates reprogramming of human induced pluripotent stem cells. Nat. Genet. 42, 1113–1117 (2010).

61. Cheng, E.-C. & Lin, H. Repressing the repressor: a lincRNA as a MicroRNA sponge in embryonic stem cell self-renewal. Developmental cell vol. 25 1–2 (2013).

62. Wang, Y. et al. Endogenous miRNA sponge lincRNA-RoR regulates Oct4, Nanog, and Sox2 in human embryonic stem cell self-renewal. Dev. Cell 25, 69–80 (2013).

63. Sridhar, B. et al. Systematic Mapping of RNA-Chromatin Interactions In Vivo. Curr. Biol. **27**, 602–609 (2017).

64. Yankova, E. et al. Small-molecule inhibition of METTL3 as a strategy against myeloid leukaemia. Nature 593, 597–601 (2021).

65. Konermann, S. et al. Transcriptome Engineering with RNA-Targeting Type VI-D CRISPR Effectors. Cell 173, 665–676.e14 (2018).

66. Yan, W. X. et al. Cas13d Is a Compact RNA-Targeting Type VI CRISPR Effector Positively Modulated by a WYL-Domain-Containing Accessory Protein. Mol. Cell 70, 327– 339.e5 (2018).

67. Zheng, G. et al. ALKBH5 is a mammalian RNA demethylase that impacts RNA metabolism and mouse fertility. Mol. Cell 49, 18–29 (2013).

68. Xia, Z. et al. Epitranscriptomic editing of the RNA N6-methyladenosine modification by dCasRx conjugated methyltransferase and demethylase. Nucleic Acids Res. 49, 7361–7374 (2021).

69. Kang, J. et al. Simultaneous deletion of the methylcytosine oxidases Tet1 and Tet3 increases transcriptome variability in early embryogenesis. Proc. Natl. Acad. Sci. U. S. A. 112, E4236–45 (2015).

70. de la Rica, L. et al. TET-dependent regulation of retrotransposable elements in mouse embryonic stem cells. Genome Biol. 17, 234 (2016).

71. Zhang, P. et al. L1 retrotransposition is activated by Ten-eleven-translocation protein 1 and repressed by methyl-CpG binding proteins. Nucleus 8, 548–562 (2017).

72. Deniz, Ö., de la Rica, L., Cheng, K. C. L., Spensberger, D. & Branco, M. R. SETDB1 prevents TET2-dependent activation of IAP retroelements in naïve embryonic stem cells. Genome Biol. 19, 6 (2018).

73. Hescheler, J. & Hofer, E. Adult and Pluripotent Stem Cells: Potential for Regenerative Medicine of the Cardiovascular System. (Springer Science & Business, 2014).

74. Verma, N. et al. TET proteins safeguard bivalent promoters from de novo methylation in human embryonic stem cells. Nat. Genet. 50, 83–95 (2018).

75. Dixon, G. et al. QSER1 protects DNA methylation valleys from de novo methylation. Science 372, (2021).

76. Liu, X. S. et al. Editing DNA Methylation in the Mammalian Genome. Cell 167, 233– 247.e17 (2016).

77. Ziller, M. J. et al. Charting a dynamic DNA methylation landscape of the human genome. Nature 500, 477–481 (2013).

78. Robertson, K. D. DNA methylation and human disease. Nat. Rev. Genet. 6, 597–610 (2005).

79. Zhang, T. et al. Active endogenous retroviral elements in human pluripotent stem cells play a role in regulating host gene expression. Nucleic Acids Res. 50, 4959–4973 (2022).

80. Takahashi, K. et al. The pluripotent stem cell-specific transcript ESRG is dispensable for human pluripotency. Cold Spring Harbor Laboratory 2020.11.25.397935 (2020) doi:10.1101/2020.11.25.397935.

81. Zhu, Z. et al. PHB Associates with the HIRA Complex to Control an Epigenetic-Metabolic Circuit in Human ESCs. Cell Stem Cell 20, 274–289.e7 (2017).

82. Sanjana, N. E., Shalem, O. & Zhang, F. Improved vectors and genome-wide libraries for CRISPR screening. Nat. Methods 11, 783–784 (2014).

83. Diao, Y., Wang, X. & Wu, Z. SOCS1, SOCS3, and PIAS1 promote myogenic differentiation by inhibiting the leukemia inhibitory factor-induced JAK1/STAT1/STAT3 pathway. Mol. Cell. Biol. 29, 5084–5093 (2009).

84. Chelmicki, T. et al. m6A RNA methylation regulates the fate of endogenous retroviruses. Nature (2021) doi:10.1038/s41586-020-03135-1.

85. Zeng, Y. et al. Refined RIP-seq protocol for epitranscriptome analysis with low input materials. PLoS Biol. 16, e2006092 (2018).

86. Picelli, S. et al. Full-length RNA-seq from single cells using Smart-seq2. Nat. Protoc. 9, 171–181 (2014).

87. Chen, S., Zhou, Y., Chen, Y. & Gu, J. fastp: an ultra-fast all-in-one FASTQ preprocessor. Bioinformatics vol. 34 i884–i890 (2018).

88. Kim, D., Paggi, J. M., Park, C., Bennett, C. & Salzberg, S. L. Graph-based genome alignment and genotyping with HISAT2 and HISAT-genotype. Nat. Biotechnol. 37, 907– 915 (2019).

89. Liao, Y., Smyth, G. K. & Shi, W. featureCounts: an efficient general purpose program for assigning sequence reads to genomic features. Bioinformatics 30, 923–930 (2014).

90. Love, M. I., Huber, W. & Anders, S. Moderated estimation of fold change and dispersion for RNA-seq data with DESeq2. Genome Biol. 15, 550 (2014).

91. Huang, D. W., Sherman, B. T. & Lempicki, R. A. Systematic and integrative analysis of large gene lists using DAVID bioinformatics resources. Nat. Protoc. 4, 44–57 (2009).

92. Sherman, B. T. et al. DAVID: a web server for functional enrichment analysis and functional annotation of gene lists (2021 update). Nucleic Acids Res. (2022) doi:10.1093/nar/gkac194.

93. Langmead, B. & Salzberg, S. L. Fast gapped-read alignment with Bowtie 2. Nat. Methods 9, 357–359 (2012).

94. Li, H. & Durbin, R. Fast and accurate short read alignment with Burrows–Wheeler transform. Bioinformatics 25, 1754–1760 (2009).

95. Li, Handsaker, Wysoker, Fennell & Ruan. The Sequence alignment/map (SAM) format and SAMtools. Bioinformatics.

96. Tarasov, A., Vilella, A. J., Cuppen, E., Nijman, I. J. & Prins, P. Sambamba: fast processing of NGS alignment formats. Bioinformatics 31, 2032–2034 (2015).

97. Feng, J., Liu, T., Qin, B., Zhang, Y. & Liu, X. S. Identifying ChIP-seq enrichment using MACS. Nat. Protoc. 7, 1728–1740 (2012).

98. Heinz, S. et al. Simple combinations of lineage-determining transcription factors prime cis-regulatory elements required for macrophage and B cell identities. Mol. Cell 38, 576–589 (2010).

99. Ramírez, F. et al. deepTools2: a next generation web server for deep-sequencing data analysis. Nucleic Acids Res. 44, W160–5 (2016).

100. Quinlan, A. R. & Hall, I. M. BEDTools: a flexible suite of utilities for comparing genomic features. Bioinformatics 26, 841–842 (2010).

